# Mapping the spatiotemporal continuum of structural connectivity development across the human connectome in youth

**DOI:** 10.1101/2024.06.17.599267

**Authors:** Xiaoyu Xu, Hang Yang, Jing Cong, Haoshu Xu, Jason Kai, Shaoling Zhao, Yang Li, Haochang Shou, Kangcheng Wang, Valerie J. Sydnor, Ting Xu, Fang-Cheng Yeh, Zaixu Cui

**Affiliations:** State Key Laboratory of Cognitive Neuroscience and Learning, Beijing Normal University; Beijing, 100875, China; Beijing Institute for Brain Research, Chinese Academy of Medical Sciences & Peking Union Medical College; Beijing, 102206, China; Chinese Institute for Brain Research, Beijing; Beijing, 102206, China; Academy for Advanced Interdisciplinary Studies, Peking University; Beijing, 100871, China; Child Mind Institute, Center for the Integrative Developmental Neuroscience; New York, NY, USA; Department of Biostatistics, Epidemiology and Informatics, Perelman School of Medicine, University of Pennsylvania; Philadelphia, PA, USA; School of Psychology, Shandong Normal University; Jinan, 250358, China; Department of Psychiatry, University of Pittsburgh Medical Center; Pittsburgh, PA, USA; Department of Neurological Surgery, University of Pittsburgh; Pittsburgh, PA, USA

**Keywords:** Adolescence, Development, Diffusion MRI, Structural Connectivity, Sensorimotor-association Connectional Axis

## Abstract

Childhood and adolescence are marked by protracted developmental remodeling of cortico-cortical structural connectivity. However, the spatiotemporal variability of white matter connectivity development across the human connectome and its relevance to cognition and psychopathology remains unclear. Using diffusion MRI data from three independent developmental cohorts spanning youth, we identified a robust divergence in structural connectivity maturation along a predefined sensorimotor-association (S-A) connectional axis during youth (http://connectcharts.cibr.ac.cn). This developmental continuum ranged from early childhood increases in sensorimotor-sensorimotor connectivity strength to late adolescent increases in association-association connectivity strength, with the transition occurring around age 15. The S-A connectional axis also captured spatial variations in the associations between structural connectivity and both higher-order cognition and general psychopathology. Moreover, group-level developmental trajectories of structural connectivity differed by cognitive and psychopathological levels, with psychopathological effects predominantly observed in association connections. These findings delineate a spatiotemporal continuum of structural connectivity development during youth, providing a normative reference for quantifying developmental variability in psychiatric disorders.

## Introduction

Myelinated axons play a central role in neuronal signal conduction, with large bundles of parallel axons comprising macroscopic white matter tracts^1^. These white matter tracts interconnect the human cerebral cortex, forming a complex network of structural connectivity (SC) known as the connectome^2^. Both animal studies and human neuroimaging have provided evidence that white matter connectivity is refined throughout childhood and adolescence^3–7^. This developmental refinement arises from microscale processes such as myelination and alterations of axon diameter, which occur during varying periods across tracts^1,5,6^. Elucidating how these changes progress spatially and temporally across the connectome can reveal how the brain prioritizes the maturation of specific connections at distinct developmental stages, and how heterogeneity in connection-specific developmental refinement impacts cognitive development. Such understanding provides insight into how SC is susceptible to influences such as exposure to psychopathology and interventions at distinct developmental periods.

Activity- and experience-dependent plasticity in myelination and axonal remodeling are major drivers of white matter connectivity maturation in youth^5,6,8,9^. During early development, sensorimotor connections experience high levels of neural activity transmission due to the rapid acquisition of motor skills and exposure to new sensory inputs. This heightened activity leads to increased expression of growth factors such as brain-derived neurotrophic factor (BDNF) and neuregulin-1, which promote axonal remodeling, dendritic arborization, and myelination in sensorimotor pathways^8^. In contrast, association connections undergo a more prolonged period of development into young adulthood, which may be attributed to continued cognitive development and the capacity for more varied experiences to engage the neural circuits underlying higher-order cognitive functions^10^. However, beyond this coarse division between sensorimotor and association connections, there is marked spatiotemporal variability in the developmental patterns of SC across the human connectome, which remains under-characterized. Moreover, how this connection-specific heterogeneity relates to individual differences in cognition and psychopathology during youth is not well understood.

Recent studies support a unifying developmental framework that cortical maturation proceeds asynchronously along a sensorimotor-association (S-A) cortical axis: a hierarchical continuum spanning from primary sensorimotor to transmodal association cortices^10–13^. This framework posits that sensorimotor cortices mature earliest, whereas association cortices exhibit protracted refinement, with a continuous spectrum between them. Here, we tested the hypothesis that the developmental maturation of white matter SC is spatiotemporally organized along the S-A axis of the human connectome, with a continuous spectrum of trajectories ranging from early-maturing sensorimotor-sensorimotor to late-maturing association-association connections. As brain development in youth is linked to both higher-order cognition and a variety of mental disorders^3,10,14^, we further hypothesized that the spatiotemporal heterogeneity in SC development would have implications for both cognition and psychopathology.

We investigated these hypotheses using diffusion magnetic resonance imaging (dMRI) tractography, which reconstructs white matter pathways by modeling water diffusion within tissues^15,16^. Leveraging the canonical S-A cortical axis^10^, we defined an S-A connectional axis that spans the full connectome from sensorimotor-sensorimotor to association-association connections. SC strength was quantified as the number of streamlines linking pairs of large-scale cortical systems, reconstructed using probabilistic tractography. We hypothesized that the developmental changes in connectivity strength would be primarily characterized by heterochronous increases that align with the S-A connectional axis, showing early strengthening in sensorimotor pathways and delayed increases in association pathways. Prior work has shown that increased structural-connectome segregation within association networks is related to better performance in higher-order cognition^3^. Therefore, we hypothesized that large-scale SC strength would decline with better cognitive performance, and these effects would exhibit spatial gradients along the S-A connectional axis, with stronger effects in higher-order association connections. Furthermore, given that mental disorders in youth are characterized by abnormal neurodevelopment^14^, we hypothesized significant associations between SC strength and psychopathological symptoms, and the strength of these associations would increase along the S-A connectional axis. To ensure the generalizability and reliability, we tested these hypotheses across three large, independent developmental datasets encompassing diverse populations. Altogether, this work establishes a connectome-wide, axis-based framework for understanding the asynchronous maturation of SC and its implications for cognition and mental health during youth.

## Results

To delineate how age-related refinements in SC spatiotemporally progress across the human connectome, we analyzed three independent developmental datasets comprising structural and diffusion MRI data. The first dataset included 590 typically developing youth aged 8.1–21.9 years from the Lifespan Human Connectome Project in Development^17^ (HCP-D; **Fig. 1a**, **Table S1**). This dataset served as the discovery sample due to its broad age range, high image quality, and harmonized acquisitions across sites. The second dataset comprised children and adolescents from the longitudinal Adolescent Brain Cognitive Development (ABCD) study^18^, including baseline data (N = 3,949, aged 8.9–11.0 years) and two-year follow-up data (N = 3,155, aged 10.6–13.8 years; **Fig. 1b**, **Table S2**). The third dataset included 947 MRI scans from Chinese youth aged 6.1–23.4 years (**Fig. 1c**, **Table S3**), enabling assessment of cross-cultural generalizability. Participant inclusion and exclusion flowcharts are provided in **Fig. S1**–**S3**.

**Fig. 1.**
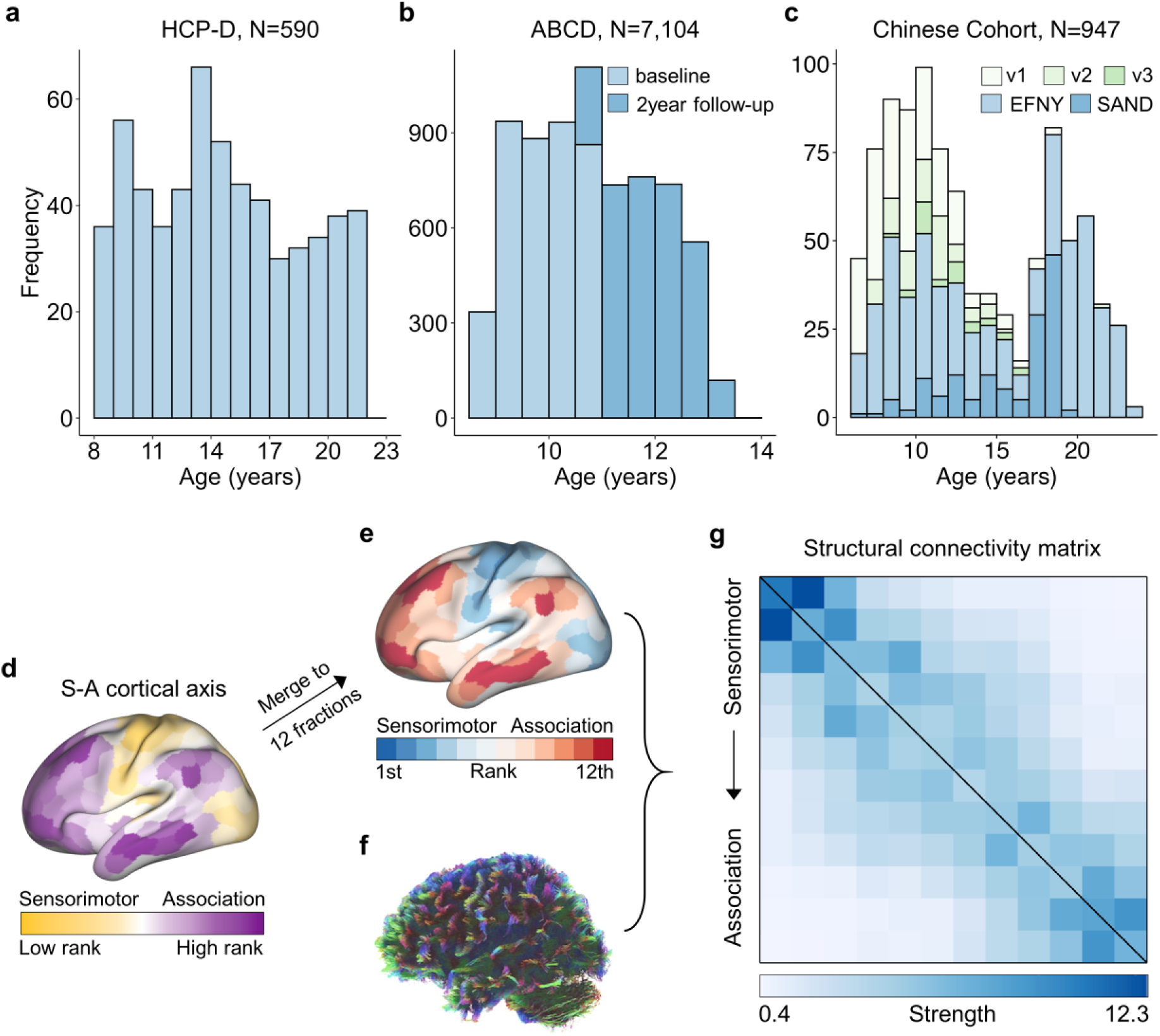
Age distribution of participants and structural connectivity construction. **a,** Age distribution (8.1–21.9 years) of 590 participants from the HCP-D dataset. **b,** Age distribution of participants from the ABCD study, including baseline (N = 3,949, 8.9–11.0 years) and 2-year follow-up (N = 3,155, 10.6–13.8 years). **c,** Age distribution of participants from the Chinese cohort, which included data from three studies: the devCCNP, EFNY, and SAND. The devCCNP is an accelerated-longitudinal dataset with up to three visits per participant. Green colors in the stacked bar chart represent the different studies and devCCNP visits. **d**, S-A cortical axis map derived by averaging multiple cortical neurobiological properties, where cortical regions are continuously ranked from lower-order sensorimotor to higher-order association cortices. **e**, Cortical atlas comprising 12 approximately equal-sized systems generated by dividing regions evenly along the S-A axis^10^. **f**, Whole-brain white matter tracts reconstructed from diffusion MRI data using probabilistic tractography. **g**, SC matrix (12 × 12) representing the number of streamlines connecting each pair of cortical systems, normalized by their respective cortical volumes. The matrix is ordered according to the S-A axis rank of the systems, from sensorimotor to association. HCP-D: the Lifespan Human Connectome Project in Development; ABCD: the Adolescent Brain Cognitive Development; devCCNP: developing Chinese Color Nest Project; EFNY: the Executive Function and Neurodevelopment in Youth; SAND: Shandong Adolescent Neuroimaging Project on Depression; v1, v2, v3: 1^st^, 2^nd^, and 3^rd^ visit from the devCCNP; S-A: sensorimotor-association.

To define individuals’ large-scale white matter connectivity, we first partitioned the cerebral cortex into 12 large-scale systems of approximately equal size along a priori defined sensorimotor-association (S-A) cortical axis (**Fig. 1d**), which was derived by averaging multiple cortical neurobiological properties^10^. These systems progressively spanned from primary sensorimotor to higher-order association cortices (**Fig. 1e**, **Fig. S4**), with each cortical system comprising regions with similar neurobiological profiles in anatomy, function, evolutionary expansion, metabolism, and gene expression. This axis-based partition was not intended to redefine canonical functional networks but to provide spatially ordered, coarse-scale cortical subdivisions for assessing how SC development varies along the S-A axis. We used 12 systems for the main analyses, representing an intermediate scale relative to the widely used Yeo-7 and Yeo-17 functional network parcellations. Robustness was assessed using alternative parcellations in which the cortex was subdivided into (i) 7 or 17 systems based on S-A axis ranks, and (ii) the canonical Yeo-7 and Yeo-17 functional network parcellations^19^.

After defining the 12 cortical systems along the S-A axis, we reconstructed individuals’ whole-brain white matter tracts (**Fig. 1f**) using probabilistic fiber tractography with multi-shell, multi-tissue constrained spherical deconvolution^20^. Anatomically constrained tractography (ACT)^21^ and spherical deconvolution informed filtering of tractograms (SIFT)^22^ were applied to improve the biological accuracy. We then quantified the number of streamlines connecting each pair of cortical systems, normalized by their respective cortical volumes, yielding a 12 × 12 SC matrix for each participant (**Fig. 1g**). These structural connectome matrices were ordered according to systems’ ranks along the S-A axis, progressing from lower-ranked sensorimotor to higher-ranked association cortices.

### Developmental refinement of structural connectivity varies across the connectome

We initially investigated the refinement of SC from ages 8.1 to 21.9 years using the HCP-D dataset^17^. Using generalized additive models (GAMs), we found that 70 out of 78 connectivity edges exhibited a significant age-related developmental effect (*Pfdr* < 0.05), while controlling for sex and head motion (**Fig. 2a**). We assessed the magnitude of age effects using the variance explained by age (partial *R*^2^) and determined their direction based on the sign of the average first derivative of the age smooth function. Our analysis revealed variations in age effects across all 78 connections, with the strongest effects observed in connections among primary sensorimotor systems (SS), moderate effects in connections among higher-order association systems (AA), and the weakest effects in connections between sensorimotor and association systems (SA, SS > AA > SA, all pairwise permutation tests, *P*_perm_ < 0.05, **Fig. S5a**). By visualizing the developmental trajectories of each connection, we observed a continuous spectrum spanning connections that display an early steep increase followed by a plateau, to those exhibiting a late increase (**Fig. 2b**).

**Fig. 2.**
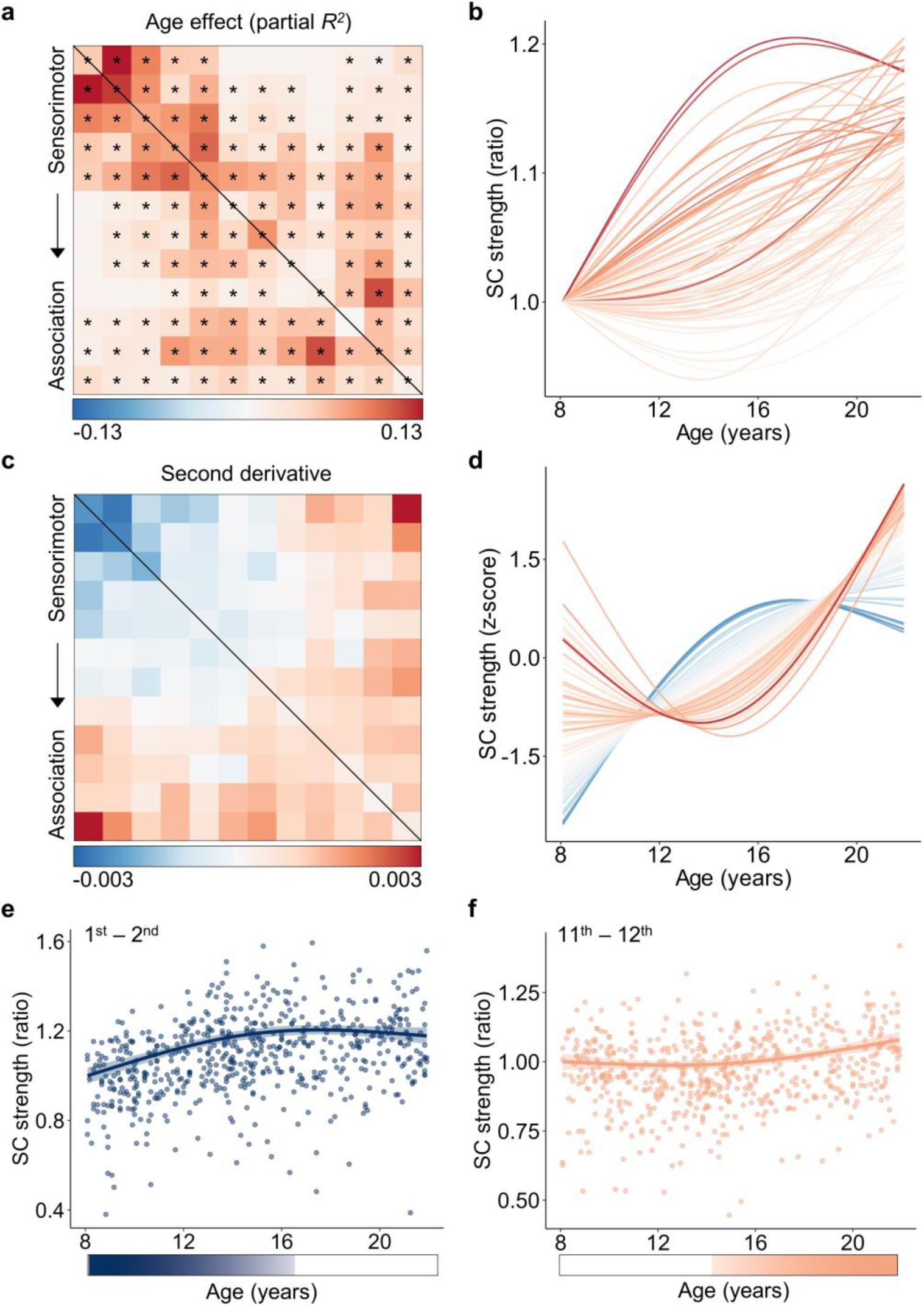
Developmental trajectories of large-scale structural connectivity vary across the connectome in youth. **a**, Age effects (partial *R*^2^) of SC strength across system-to-system connections, modeled using generalized additive models (GAMs). Black asterisks indicate significant age effects (*Pfdr* < 0.05). **b**, Developmental trajectories of SC strength showing a continuous spectrum from early increases that plateaus to later, prolonged increases. Curve colors correspond to the effect-size matrix in panel (**a**). **c**, Second derivatives of age fits revealing a continuous spectrum of developmental trajectories across connectome edges. **d**, Z-scored trajectories highlighting a continuous shift in curvature from early-maturing sensorimotor to late-maturing association connections. Curve colors correspond to the second-derivative matrix in panel (**c**). **e**, **f**, Representative developmental trajectories of SC between the 1^st^ and 2^nd^ systems involving primary visual and somatomotor cortices (**e**), and between the 11^th^ and 12^th^ systems involving higher-order frontal and temporal cortices (**f**). Each point represents one participant (N = 590); bold lines indicate GAM fits, and shaded envelope denotes 95% confidence intervals. Color bars beneath each plot mark age windows showing significant changes in SC strength, shaded by the rate of change. SC: structural connectivity.

To capture variations in the shape of these developmental trajectories, we calculated the average second derivative of the age fits for each connection, quantifying the curvature of the developmental curve (**Fig. 2c**). Negative values indicate concave-downward trajectories characterized by an earlier strengthening followed by plateaus, while positive values reflect concave-upward trajectories with delayed developmental strengthening. We observed that the second derivatives displayed a substantial heterogeneity across the connectome, with positive values predominantly in AA connections and negative values in SS connections (SS < 0, AA > 0, *P*_perm_ < 0.001, **Fig. S5b**). Z-scoring the developmental fits for each connection revealed a continuous spectrum ranging from downward-concave sensorimotor to upward-concave association connections (**Fig. 2d**). Illustrative examples highlight this temporal gradient: connectivity between primary sensorimotor systems showed pronounced strengthening during childhood, peaking around mid-adolescence (**Fig. 2e**), whereas association-association connectivity remained relatively stable through early adolescence before accelerating into young adulthood (**Fig. 2f**). Assessing the rate of developmental change within age windows confirmed that the declines were not significant for sensorimotor connection during late period (**Fig. 2e**) or for association connection during early period (**Fig. 2f**). Across all 78 connections, no significant (*P_FDR_* < 0.05) decreases in connectivity strength were observed at any age (**Fig. S6**).

### Developmental variability of structural connectivity aligns with the S-A connectional axis

Having established that SC developmental trajectories exhibit marked heterogeneity across the connectome, we next evaluated whether this variability spatially aligns with the S-A axis. The S-A cortical axis serves as a unifying cortical organizing principle encompassing diverse neurobiological properties, with a continuous progression observed along the axis from unimodal sensorimotor to transmodal association cortices^10^. To compare edge-level developmental variability to this hierarchical axis, we derived an S-A connectional axis. Each of the 78 system-to-system connections was assigned an S-A rank by summing the squared cortical S-A ranks of the two connected systems. This procedure yields higher connectional ranks for links involving higher-order association regions and lower ranks for connections among sensorimotor systems (**Fig. 3a**). This approach is motivated by evolutionary principles, as transmodal association areas are among the most recently expanded in primate evolution and support integrative cognitive functions. Accordingly, connections involving these regions likely represent more evolutionarily advanced and functionally integrative pathways. The S-A connectional axis was highly correlated with both the functional gradient^23^-based connectional axis (rho = 0.98; **Fig. S7**) and the T1w/T2w^24^-based connectional axis (rho = −0.90; **Fig. S7**), supporting the S-A connectional axis as an integrative representation of hierarchical cortical organization.

**Fig. 3.**
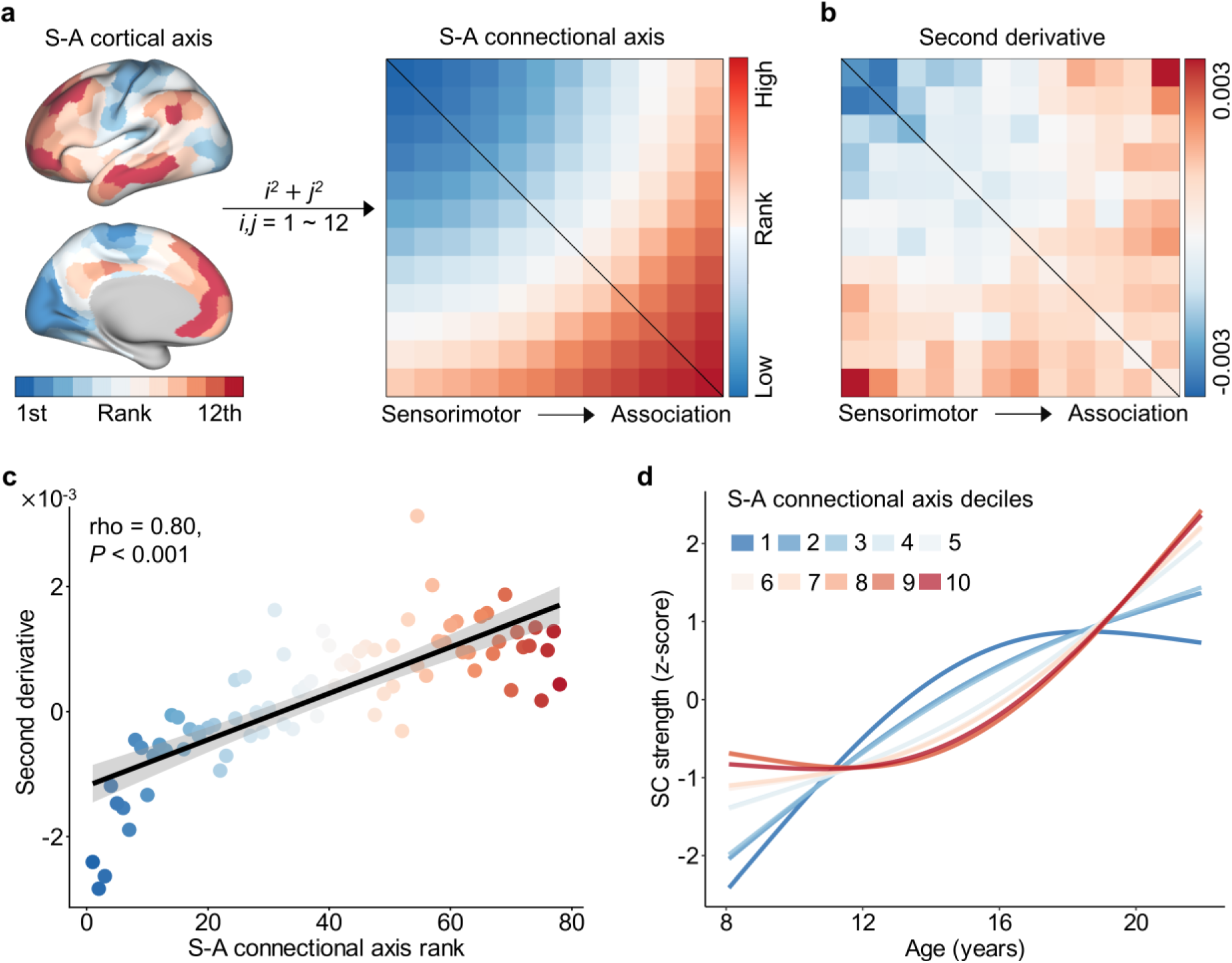
The heterogeneity of structural connectivity development across the connectome aligns with the S-A connectional axis. **a**, Construction of the S-A connectional axis derived from the priori S-A cortical axis. A single connectional rank was assigned to each connection by summing the squared S-A cortical ranks of the two connected systems. These connectional ranks were scaled into discrete values from 1 to 78, with lower ranks reflecting sensorimotor and higher ranks reflecting association connections. **b**, Second derivatives of developmental trajectories in SC strength (from Fig. 2c). **c**, Significant positive correlation between the second derivative and S-A connectional axis rank across all connections (rho = 0.80, *P* < 0.001). Point colors correspond to the connectional axis matrix in (a). **d**, Averaged developmental trajectories of SC strength across deciles of the S-A connectional axis. The connectional axis was divided into 10 bins, each consisting of 7–8 connections; age fits were averaged within each bin and z-scored for visualization. The first decile (dark blue) represents the sensorimotor pole and the tenth decile (dark red) represents the association pole of the axis. Maturation trajectories diverged most between the two poles and varied continuously between them. SC: structural connectivity; S-A: sensorimotor-association.

We next evaluated whether developmental curvatures defined by the mean second derivative from the GAM fits (**Fig. 3b**, also see **Fig. 2c**) varies systematically along the S-A connectional axis. Spearman’s rank correlation revealed a strong positive association between the second derivatives and S-A connectional axis rank (rho = 0.80, *P* < 0.001, **Fig. 3c**). Connections near the sensorimotor pole exhibited negative second derivatives, indicating early, concave-downward maturation, whereas connections closer to the association pole displayed positive derivatives, reflecting later, concave-upward trajectories. Intermediate connections fell along a continuous spectrum between these two extremes. To further illustrate these patterns, we grouped the 78 connections into deciles based on their S-A connectional axis rank and averaged the developmental trajectories within each bin. This visualization confirmed a graded progression of maturation along the axis (**Fig. 3d**): connections at the sensorimotor end showed early strengthening followed by plateau, whereas those at the association end exhibited delayed but prolonged increases across adolescence.

### Developmental alignment with the S-A connectional axis shifts during youth

Having shown that SC developmental heterogeneity aligns with the S-A connectional axis, we further evaluated how this alignment evolves throughout youth. We divided the age range of 8.1–21.9 years into 1,000 equally spaced intervals and estimated the developmental rate within each segment. Given the brevity of the interval, we assumed a linear developmental effect in this short period and quantified the rate with the first derivative of age-related change. Visualizing these age-resolved change rates across the S-A connectional axis (**Fig. 4a**) revealed a continuous spatiotemporal progression: early and mid-childhood were dominated by positive developmental rates in sensorimotor connections with lower ranks, whereas late adolescence and early adulthood showed positive rates concentrated in association connections with higher ranks. No significant negative change rates were observed for any connection (**Fig. S6**).

**Fig. 4.**
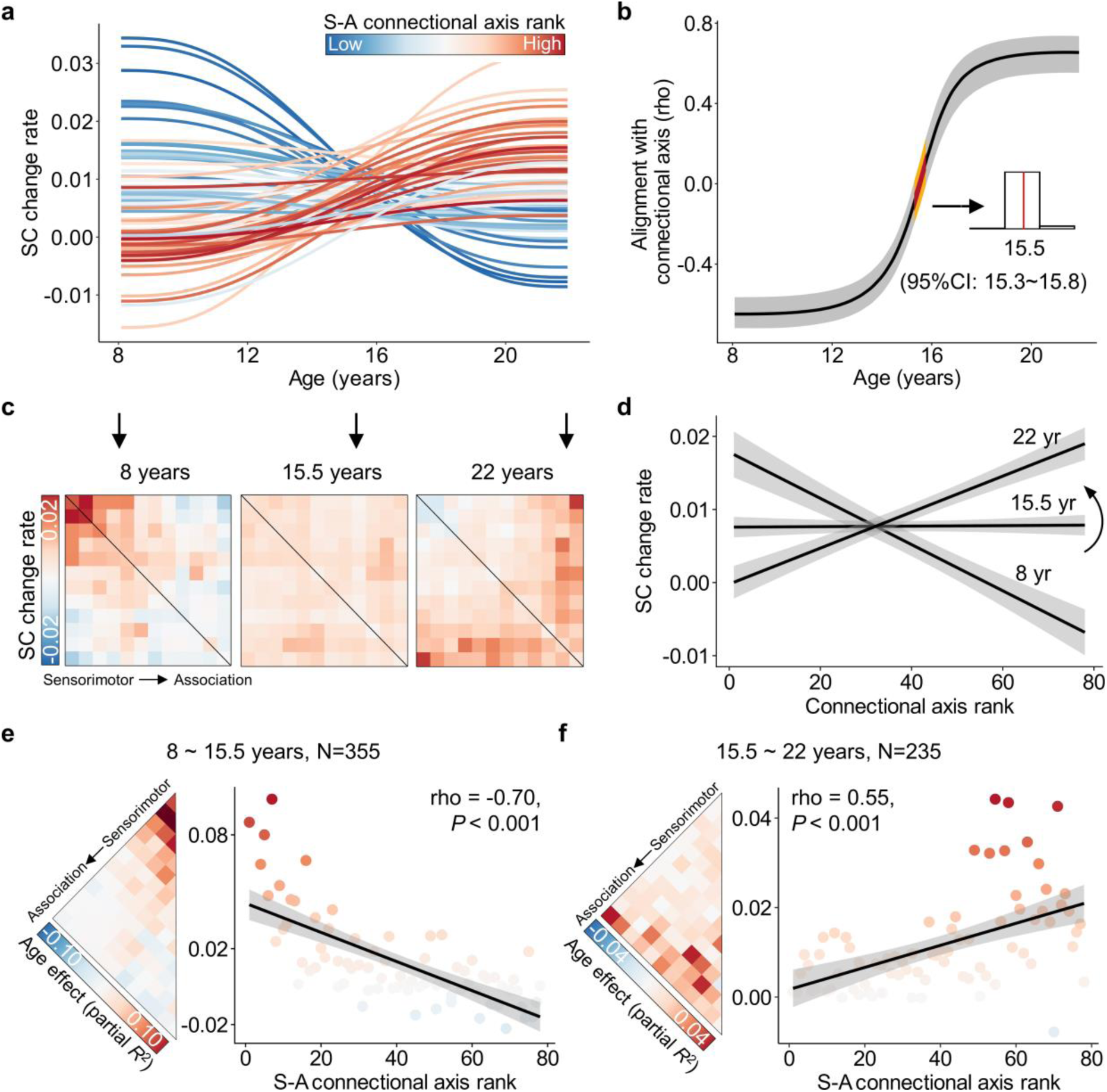
The spatial alignment between structural connectivity development and the S-A connectional axis shifts throughout youth. **a,** Developmental rates (first derivatives) of large-scale structural connections from ages 8.1 to 21.9 years. Each line represents one connection, color-coded by its S-A connectional axis rank. Connections at the sensorimotor pole of the S-A connectional axis show high positive change rates in childhood that decline toward adulthood, whereas those at the association pole display the opposite pattern. A continuous spectrum of intermediate developmental patterns spans the two poles. **b**, Age-resolved alignment between the spatial pattern of SC developmental rates and the S-A connectional axis. Alignment increases from a strong negative association in childhood to a strong positive association in young adulthood, crossing zero around 15.5 years. To estimate uncertainty, we drew 1,000 samples from the posterior derivatives of each connection’s age smooth function and then evaluate age-resolved correlations between these derivatives and S-A connectional axis ranks for each sample. The black line indicates the median correlation, the gray band indicates the 95% CI, and the yellow band marks the 95% CI for the age of zero correlation. The inset histogram shows the distribution of zero-alignment ages (15.3–15.8 years; median 15.5). **c**, Matrices of age-specific developmental change rates (first derivatives) at ages 8.1, 15.5, and 21.9 years. **d**, Scatterplots showing age-specific alignments between developmental change rates and S-A connectional axis ranks across all edges, illustrating the continuous transition from negative to positive alignment. **e**, **f**, Divergent developmental patterns of SC between younger and older youths. Age effects (partial *R*^2^) show a negative correlation with S-A connectional axis ranks in younger youths (**e**, 8.1–15.5 years; rho = −0.70, *P* < 0.001) and a positive correlation in older youths (**f**, 15.5–21.9 years; rho = 0.55, *P* < 0.001). Point colors correspond to age effects in the left matrix; two outlier edges (effect sizes = 0.18 and 0.16) were excluded from panel (**e**). SC: structural connectivity; S-A: sensorimotor-association; CI: credible interval.

We next evaluated how the spatial alignment between SC developmental rates and S-A connectional axis ranks evolves from age. To achieve this, we calculated the age-resolved spatial alignment between the connectome-wide change rates and S-A connectional axis for each of 1,000 age spaced intervals. This analysis revealed a continuous shift from negative to positive alignment across development (**Fig. 4b**). The alignment was stable before ∼13 years, increased rapidly during mid-adolescence, and stabilized in young adulthood. The alignment crossed zero at a median age of 15.5 years (95% credible interval, CI, 15.3–15.8), indicating a group-level shift in the spatial organization of connectivity strengthening. Early in childhood, developmental increases were strongest at the sensorimotor pole and decreased along the S-A axis (**Fig. 4c**, left). After mid-adolescence, the pattern inverted with stronger increases emerged toward the association pole (**Fig. 4c**, right). Matrices of developmental change rates at ages 8.1, 15.5, and 21.9 years illustrated this transition from anti-aligned to aligned with the S-A connectional axis (**Fig. 4c**), a shift confirmed by scatterplots of change rates versus S-A ranks (**Fig. 4d**).

Given the inflection point at 15.5 years, we next compared developmental effects before and after this transition. Participants were divided into two subgroups (8.1–15.5 years, N = 355; 15.5–21.9 years, N = 235), and age effects were re-estimated within each group using GAM while controlling for sex and head motion. Before 15.5 years, age effects were strongest in sensorimotor connections and weaker or negative in association connections (**Fig. 4e**, left), yielding a significant negative correlation with S-A connectional axis (rho = −0.70, *P* < 0.001, **Fig. 4e**, right). After 15.5 years, the pattern reversed: association connections exhibited greater age effects than sensorimotor connections (**Fig. 4f**, left), resulting in a significant positive correlation with S-A connectional axis (rho = 0.55, *P* < 0.001, **Fig. 4f, right**). Together, these findings indicate a rapid mid-adolescent shift in the developmental program of SC maturation, with connectivity strengthening progressively transitioning from sensorimotor to association systems after approximately 15.5 years of age. Considering that this period overlaps with puberty, we further examined whether the timing of this shift differed by sex. We fitted developmental models for each of the 78 connections separately in females (N = 317; **Fig. S8a**) and males (N = 273; **Fig. S8b**), and estimated the age at which the spatial alignment between developmental rates and the S-A axis crossed zero in each group. This zero-crossing age point, marking the shift from sensorimotor-dominant to association-dominant refinement, occurred slightly earlier in females (15.1 years, 95% CI: 14.8–15.5; **Fig. S8c**) than in males (16.2 years, 95% CI: 15.7–16.7; **Fig. S8d**). Participant-level permutation testing indicated that this sex difference was statistically significant (1,000 permutations, *P*_perm_ = 0.005; **Fig. S8e**). Together, these findings suggest that the timing of the shift in SC development from sensorimotor-dominant to association-dominant is modestly advanced in females relative to males.

### Generalizability and robustness of developmental variability along the S-A connectional axis

To assess whether the alignment between SC development and the S-A connectional axis generalizes across datasets and remains robust to methodological variations, we performed a series of sensitivity analyses. First, we tested generalizability of our findings using the Chinese youth cohort (ages 6.1–23.4 years; **Fig. 1c**). Significant developmental effects were observed in 66 of 78 connections (**Fig. 5a**), and the developmental trajectories were highly similar to those in the HCP-D dataset (**Fig. 5b, c**). Consistently, the average second derivatives were positively correlated with S-A connectional axis ranks (rho = 0.75, *P* < 0.001, **Fig. 5d**), and the age-resolved alignment shifted from negative to positive values, with a zero-alignment age of 15.5 years with 95% CI from 15.2 to 15.8 years (**Fig. 5e, f**). These findings also generalized to the ABCD dataset (ages 8.9–13.8 years; **Fig. 1b**), an independent cohort with a longitudinal design and a younger age range than the other two datasets. Consistent with results in the younger subset of the HCP-D sample, developmental effects on SC strength were negatively correlated with S-A connectional axis ranks (rho = −0.69, *P* < 0.001; **Fig 5g, h**), and the alignment between developmental rates and S-A ranks increased from strongly negative values toward zero across the observed age range (**Fig. 5i**). We further examined whether this S-A axis-structured developmental pattern could be detected at the within-person level. Using longitudinal data and linear mixed-effects models, we found that within-person SC age effects were also negatively correlated with S-A axis ranks across connections (rho = −0.57, *P* < 0.001; **Fig. S9a,b**).

**Fig. 5.**
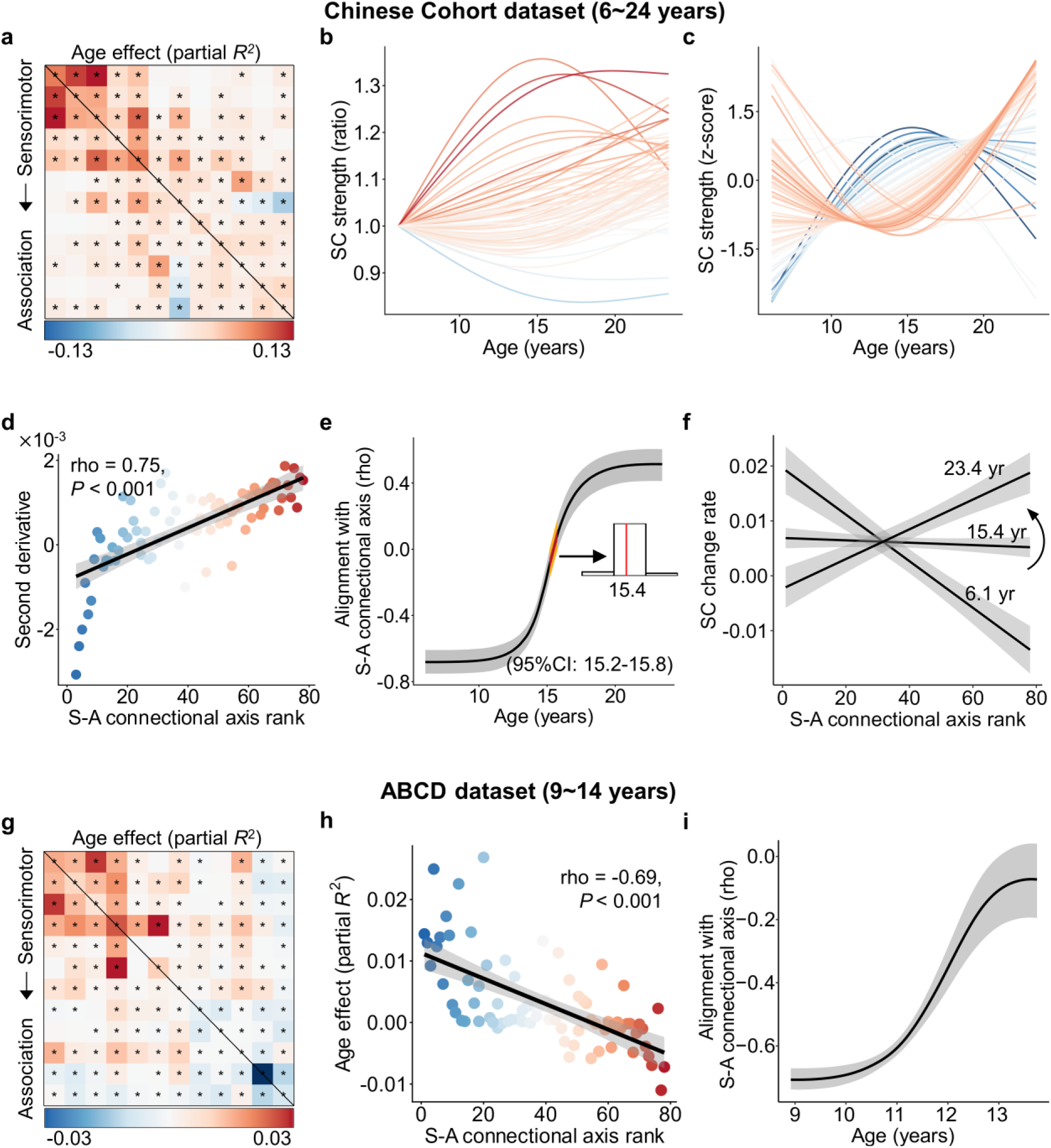
Structural connectivity development unfolds along the S-A connectional axis in two independent youth samples. Chinese Cohort dataset: **a**, Age effects (partial *R*^2^) of SC strength across system-to-system connections, with black asterisks indicating significant age effects (*P*_FDR_ < 0.05). **b**, Developmental trajectories of SC strength across the connectome, color-coded according to the effect-size matrix in a. **c**, Second derivatives of the developmental trajectories. **d**, Second derivatives were positively correlated with S-A connectional axis ranks (rho = 0.75, *P* < 0.001). **e**, Age-resolved alignment between the spatial pattern of SC developmental change rates and the S-A connectional axis across youth, with zero alignment occurring around 15.4 years (95% CI, 15.2–15.8). **f**, Age-specific relationships between SC change rates and S-A ranks at 6.1, 15.4, and 23.4 years. **ABCD dataset: g**, Age effects of SC strength across system-to-system connections. **h**, Age effects were negatively correlated with S-A connectional axis ranks across all connections (rho = −0.69, *P* < 0.001). **i**, Age-resolved alignment between the spatial pattern of SC developmental change rates and the S-A connectional axis increased from strongly negative toward near-zero values across the observed age range. SC: structural connectivity; S-A: sensorimotor-association; ABCD: the Adolescent Brain Cognitive Development.

We next evaluated the robustness of our findings within the HCP-D dataset using seven complementary methodological tests: (1) varying the number of S-A cortical systems (7 and 17 instead of 12, **Fig. S10a**,**b**); (2) reconstructing structural connectomes using the canonical Yeo-7 and Yeo-17 cortical parcellations (**Fig. 6a**,**b**); (3) regressing out Euclidean distance between system pairs when assessing S-A alignment (**Fig. 6c**); (4) controlling for mean whole-brain SC strength (**Fig. 6d**); (5) including socioeconomic status (SES; **Fig. 6e**) and intracranial volume (ICV; **Fig. 6f**) as additional covariates; (6) reconstructing connectomes from major bundle-based TractSeg^25^ tractography (**Fig. 6g**); and (7) defining an alternative S-A connectional axis based on the product of system ranks (**Fig. 6h**, **Fig. S10c**). Across these analyses, average second derivatives consistently correlated with S-A connectional axis ranks, and age-resolved alignment exhibited a reproducible transition from negative to positive, with zero alignment occurring near mid-adolescence (∼15 years). Moreover, we tested whether the S-A connectional axis captures developmental variability when the developmental axis is defined in a data-driven manner. We derived a dominant developmental axis by applying principal component analysis (PCA) to trajectories of SC developmental rates (**Fig. S11a**) and compared it with three connectional axes. The S-A connectional axis best explained the data-driven axis, showing a stronger correspondence (rho = 0.80, *P* < 0.001; **Fig. S11b**) than alternative organization principles, including the functional gradient^23^-based connectional axis (rho = 0.76, *P* < 0.001; **Fig. S11c**) and the T1w/T2w^24^-based connectional axis (rho = −0.57, *P* < 0.001; **Fig. S11d**).

**Fig. 6.**
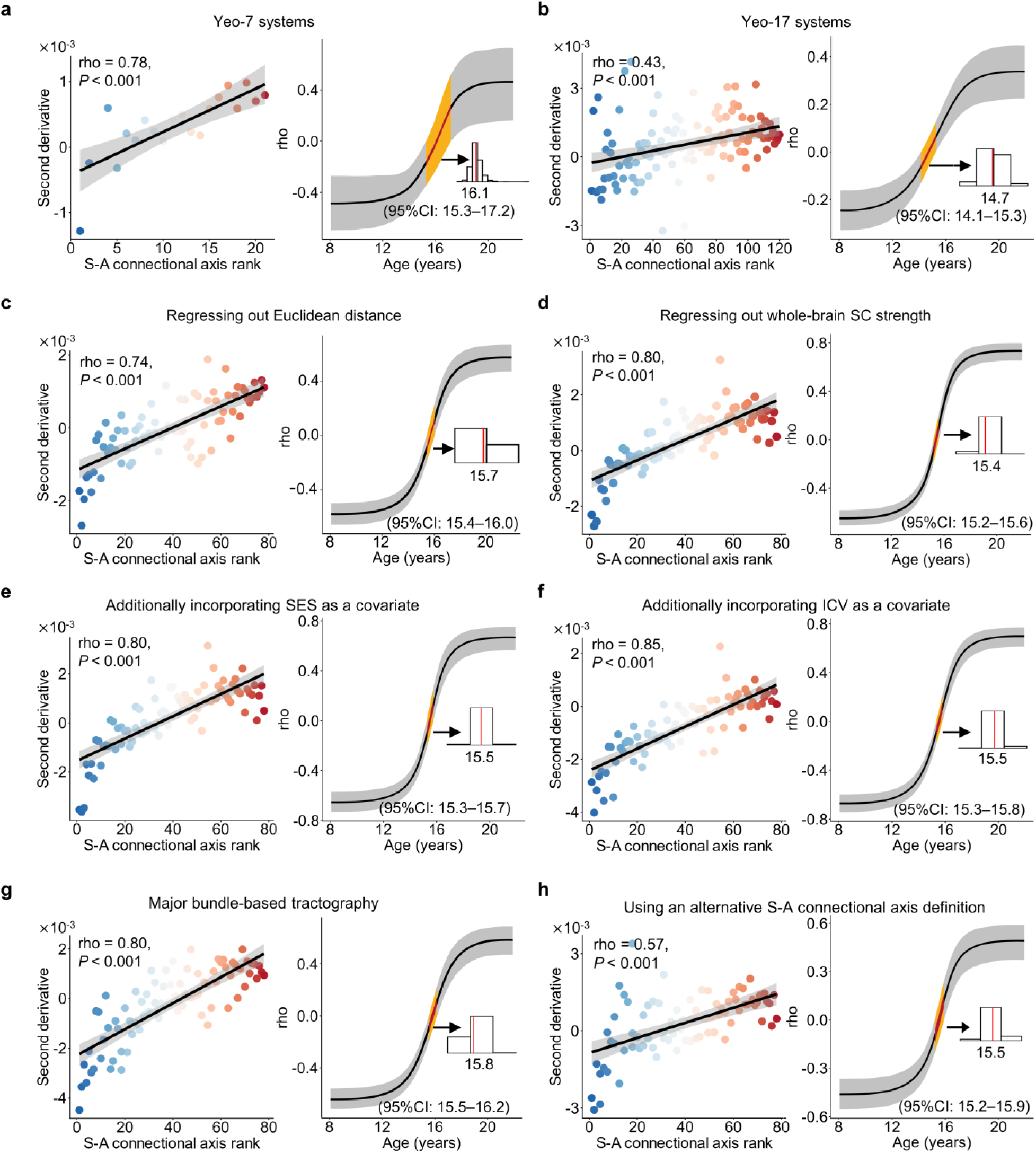
Developmental variability of structural connectivity is robust to methodological variations. We validated our results using several alternative methodological approaches, including reconstructed structural connectomes based on the canonical Yeo-7 (**a**) and Yeo-17 (**b**) cortical parcellations^19^; re-estimation of alignments after regressing out Euclidean distance between system pairs (**c**); controlling for mean whole-brain SC strength when fitting developmental models (**d**); developmental models incorporating SES (**e**) or ICV (**f**) as additional covariates; major bundle-based TractSeg tractography (**g**); and defining an alternative S-A connectional axis based on the product of system ranks (**h**). **a**–**h**, across all methodological replications using the HCP-D dataset, the primary findings were consistently reproduced. In each analysis, the left panel shows the correlation between the second derivatives of developmental trajectories and S-A connectional axis ranks, and the right panel shows the age-resolved alignment between developmental change rates and the S-A axis. Dots are color-coded by S-A ranks. The black line represents the median correlation value, and the gray band indicates the 95% CI. Yellow bands denote CI for the age of zero alignment, with the median and 95% CI of the transition age annotated. SC: structural connectivity; S-A: sensorimotor-association; SES: socioeconomic status; ICV: intracranial volume; CI: credible interval.

Overall, these sensitivity analyses demonstrate that the developmental variability of SC along the S-A connectional axis is robust to a wide range of methodological choices and generalizes across cohorts.

### Cognitive and psychopathological associations of structural connectivity vary along the S-A connectional axis

Prior work has linked SC to higher-order cognition in youth^3,4^, but the connectome-wide organization of these associations remains unclear. Here, we tested whether SC–cognition associations vary systematically along the S-A connectional axis. Using datasets from both HCP-D and ABCD, we selected the fluid composite score from the NIH Toolbox, derived from five tasks assessing working memory, inhibition, cognitive flexibility, episodic memory, and processing speed, as a measure of higher-order cognitive performance^26^ (**Fig. 7a**). We modeled associations between SC strength and cognition using GAMs controlling for the smooth term of age, sex, and head motion. Because the ABCD two-year follow-up lacked flexibility and working-memory measures, cognitive analyses were conducted using baseline data (ages 8.9–11.0 years).

**Fig. 7.**
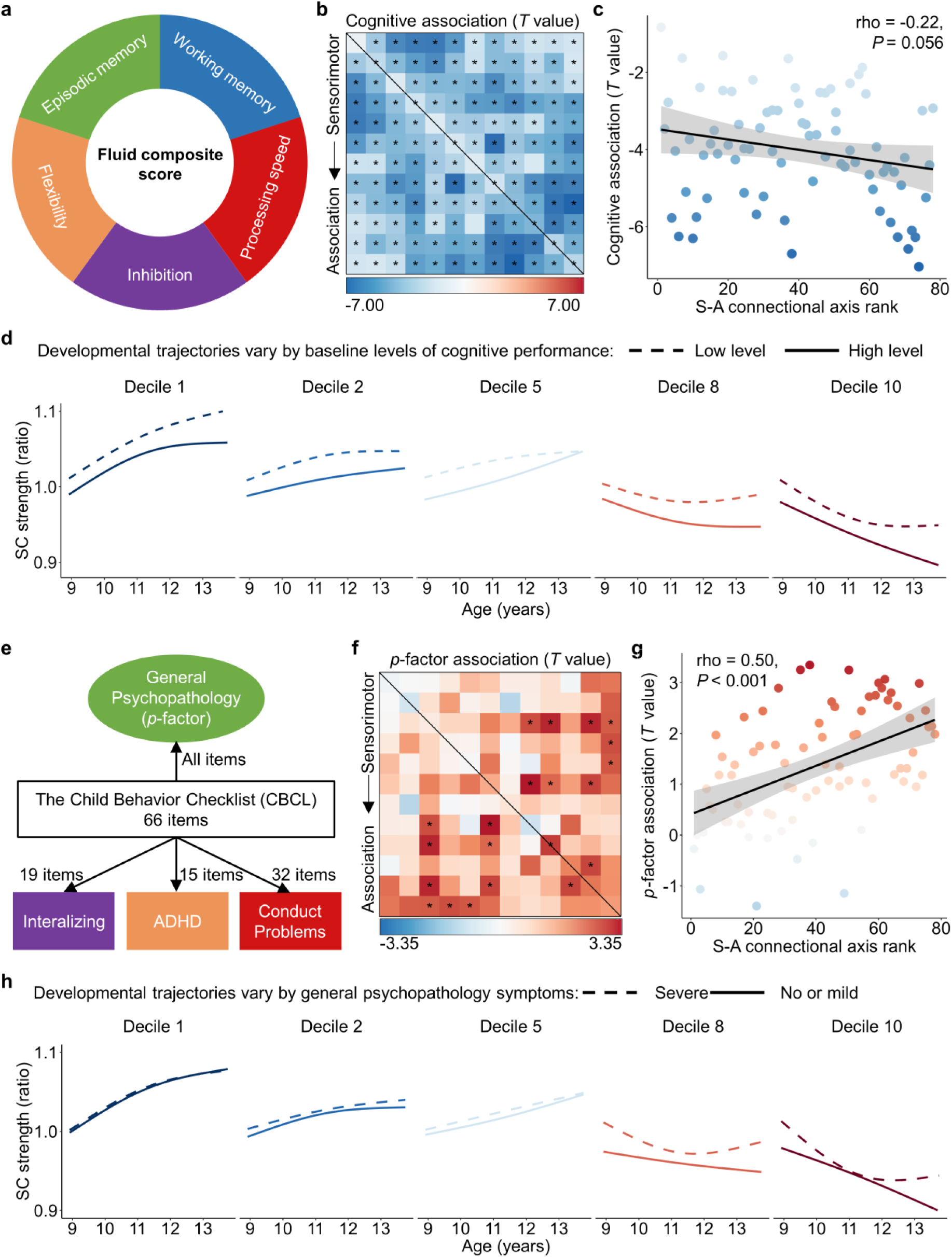
S-A connectional axis captures spatial variability in the associations between structural connectivity and both higher-order cognition and general psychopathology. **a**–**d**, Associations between large-scale SC and cognitive performance. **a**, The fluid composite score from the NIH Toolbox, derived from performance across five cognitive tasks, including working memory, inhibition, cognitive flexibility, episodic memory, and processing speed, was used to represent higher-order cognitive performance. **b**, Spatial map of associations between SC strength and cognitive performance (*T* values). Darker blue indicates stronger negative associations. Black asterisks indicate statistically significant associations (*P_FDR_* < 0.05). **c**, Across all connections, the effect sizes of the association between connectivity strength and the cognitive performance are related to S-A connectional axis ranks (rho = −0.22, *P* = 0.056), suggesting relatively stronger negative effects in higher-ranked association connections. **d**, Developmental trajectories of SC strength are shown for participants with low (10^th^ percentile) and high (90^th^ percentile) cognitive performance, across five deciles of the S-A connectional axis. **e**–**h**, Associations between SC and a general psychopathology factor (‘*p* factor’). **e**, A confirmatory bifactor analysis identified four independent psychopathology dimensions from 66 clinical items assessed by the Child Behavior Checklist (CBCL). The ‘*p*-factor’ reflects shared vulnerability to a broad range of psychiatric symptoms. **f**, Spatial map of associations between SC strength and the *p*-factor (*T* values). Darker red indicates stronger positive associations. Black asterisks indicate statistically significant associations (*P_FDR_* < 0.05). **g**, Across all connections, *p*-factor associations were positively correlated with S-A connectional axis ranks (rho = 0.50, *P* < 0.001), indicating stronger positive associations in higher-ranked association connections. **h**, Developmental trajectories of SC strength for participants with severe (90^th^ percentile) and mild or no (10^th^ percentile) psychopathology symptoms, across five S-A deciles. SC: structural connectivity; S-A: sensorimotor-association.

In the ABCD dataset, 73 of 78 connections showed significant associations between SC strength and cognition (*Pfdr* < 0.05; **Fig. 7b**; partial correlation coefficients in **Fig. S12a**). All effects were negative, indicating that weaker large-scale connectivity related to better cognitive performance. Effect sizes increased along the S-A connectional axis, with stronger effects in connections involving association systems. A connectome-wide analysis confirmed a trend toward more negative effects at higher S-A ranks (rho = −0.22, *P* = 0.056; **Fig. 7c**), suggesting a trend-level correlation between SC–cognition associations and the S-A axis ranks. To examine whether developmental trajectories differed by cognitive level, we modeled age-related changes in SC strength using generalized additive mixed models (GAMMs) with an age-by-cognition interaction, controlling for sex, head motion and participant random intercepts. High and low cognitive levels were defined by the 90^th^ and 10^th^ percentiles of baseline cognitive performance, respectively. The developmental trajectories of sensorimotor connections (deciles 1 and 2) were steeper at lower cognitive scores, whereas those of association connections (deciles 8 and 10) showed a slower decline (**Fig. 7d**). The spatial alignment of SC–cognition associations with the S-A connectional axis remained robust, and the trajectory patterns across cognitive levels remained consistent when using age-corrected fluid composite scores (**Fig. S13a,b**). In contrast, no significant SC–cognition associations were observed in the HCP-D dataset, likely reflecting limited power (N = 422) and reduced behavioral variability in this typically developing sample.

Psychiatric symptoms also emerge from neurodevelopmental perturbations, particularly within higher-order association cortex^10,14^. We therefore examined whether associations between SC and psychopathology during youth followed the S-A connectional axis. Using the ABCD dataset, we derived a general psychopathology factor (*p*-factor) from 66 Child Behavior Checklist (CBCL) items via confirmatory bifactor analysis (**Fig. 7e**), following established hierarchical model^27^. Higher *p*-factor scores reflect greater vulnerability to transdiagnostic psychiatric symptoms^28,29^.

We then modeled SC–*p*-factor associations with GAMMs including per-participant random intercepts, while controlling for age, sex, and head motion. Eleven connections showed significant positive associations (*P_FDR_* < 0.05), indicating greater SC strength with a higher *p*-factor score (**Fig. 7f**; partial correlation coefficients in **Fig. S12b**). Across the connectome, *p*-factor effects correlated positively with S-A connectional axis ranks (rho = 0.50, *P* < 0.001, **Fig. 7g**), revealing that associations between SC and psychopathology were spatially organized along the S-A connectional axis. Finally, we tested whether developmental trajectories differed by psychopathology level. SC trajectories were modeled for participants at the 10^th^ (no/mild symptoms) and 90^th^ (severe symptoms) percentiles of the *p*-factor. Differences were most pronounced near the association pole (deciles 8 and 10), paralleling the cognitive results (**Fig. 7h**). Higher *p*-factor was associated with an earlier inflection and altered strengthening in connections near the association end of the S-A axis at the group-level. Notably, the observed differences in developmental trajectories by cognition and psychopathology reflect a combination of between-person and within-person age-related variance. Detecting comparable nonlinear within-person differences will require denser longitudinal sampling. The alignment of the spatial pattern of SC–psychopathology associations with the S-A connectional axis remained robust when psychopathological burden was indexed using the CBCL Total Problems score (**Fig. S13c**). In addition, stratifying SC trajectories by CBCL Total Problems levels revealed decile-resolved developmental trajectory differences similar to those observed with the *p*-factor (**Fig. S13d**). Together, these results indicate that both the spatial organization of SC–psychopathology associations and the associated developmental patterns are robust to the operationalization of psychopathology.

Together, these findings demonstrated that both cognitive and psychopathological associations of SC are systematically patterned along the S-A connectional axis and exhibit distinct developmental trajectories across youth.

## Discussion

In this study, we delineated how SC maturation unfolds spatiotemporally across the human connectome during youth. Across three independent, racially diverse datasets, we demonstrated that developmental trajectories of SC strength systematically vary along a predefined S-A connectional axis. Particularly, we observed a continuous spectrum ranging from early age-related increases in connectivity strength between sensorimotor regions to post-adolescence increases in association-association connections at the top end of the S-A axis. The spatial alignment between connectome-wide developmental change rates and the S-A connectional axis evolved during youth, with a transition from negative to positive alignment around 15 years of age. Additionally, we found that the S-A connectional axis captures connectome-wide spatial variation in relationships between SC strength and both cognitive performance and transdiagnostic psychiatric symptomatology. Together, these results provided evidence of hierarchical development of SC along a macroscale connectional axis of the human connectome during youth.

White matter connectivity primarily consists of bundles of myelinated or unmyelinated axons connecting different brain regions^1^. Using diffusion MRI tractography, we observed sustained increases in white matter connectivity strength throughout childhood, adolescence, and young adulthood, which is consistent with prior reports of progressive white matter volume growth throughout early life that peaks at 28.7 years of age^7^. Moreover, we observed substantial heterogeneity in periods of increase across connectome edges. Connection strength began to increase before the earliest age studied (i.e., 8.1 years old) and ceased around 16 years old for unimodal sensorimotor-sensorimotor connections. In contrast, association-association connections began to strengthen around 14 years of age and exhibited rapid increases until the oldest age studied (i.e., 21.9 years old). Connections linking intermediate systems followed an orderly progression between these two extremes, forming a continuous developmental spectrum across the connectome.

These findings align with prior work showing asynchronous maturation across white matter pathways, where projection and commissural tracts mature earlier than long-range association fibers and posterior regions preceding anterior counterparts^30,31^. Our results extend these coarse observations by providing a connectome-wide, quantitative description of developmental sequencing across all cortico-cortical connections. We further showed that this spatial variation in SC development closely follows a predefined S-A connectional axis spanning from sensorimotor-sensorimotor to association-association connections. This continuum mirrors prior evidence of asynchronous cortical maturation along the S-A cortical axis, including changes in gray matter morphology^7,32^, intracortical myelin^11^, intrinsic activity^12^, and functional connectivity strength^13,33^. Together, these findings support the view that white and gray matter mature in a coordinated, axis-structured manner. An early-established white-matter scaffold^34,35^ may provide anatomical constraints on cortical morphometric development^36^, whereas white matter continues to undergo protracted refinement, such as ongoing myelination and activity-dependent plasticity^6^, into adulthood. Our results extend this axis-structured maturation framework to the structural connectome and provide initial evidence that the developmental sequence of SC conforms to the S-A axis across the human connectome. Notably, this alignment between developmental change and the S-A axis shifted from negative to positive around mid-adolescence, marking a transition from predominant strengthening of sensorimotor connections to accelerated maturation of association connections. This pattern aligns with prior reports that cortical plasticity peaks in association cortices between ages 14 and 16 before declining^12^, suggesting that the late strengthening of association connectivity may promote the functional stabilization of higher-order systems.

The hierarchical progression of white matter maturation likely reflects the interplay of molecular, cellular, and activity-dependent processes. The myelination of axonal tracts has also been demonstrated to follow a chronologic sequence, wherein fibers belonging to specific functional systems mature simultaneously^6^. Sensorimotor pathways undergo early myelination during development, whereas association pathways myelinate later during adolescence^6^. This maturational sequence in myelination is driven by oligodendrocytes and regulated by various cellular and molecular mechanisms, such as transcription factors (Olig family), growth factors (BDNF, neuregulin-1), and hormones (T3)^37^. The process of myelination is strongly influenced by experience- and activity-dependent plasticity mechanisms^8,9^. For example, studies have demonstrated that signaling molecules regulated by action potential firing in axons can impact the development of myelinating glia^8^. It is widely acknowledged that early development is primarily marked by new sensorimotor experience, whereas later developmental stages are characterized by prolonged exposure to increasingly complex cognitive and social experiences. Therefore, experience-driven and activity-dependent myelin plasticity could be a primary mechanism underlying the maturational sequence of strengthening in white matter SC.

Our findings also revealed that SC strength was associated with higher-order cognitive performance, and that SC–cognition associations showed a trend toward systematic variation along the S-A connectional axis. Across the connectome, SC strength correlated negatively with a composite measure of higher-order cognition encompassing working memory, inhibition, flexibility, episodic memory, and processing speed^38^, indicating that weaker large-scale connectivity relates to better cognitive performance. This pattern aligns with prior evidence that greater segregation among brain networks supports improved executive function^3,33,39,40^. Moreover, the magnitude of these negative associations increased along the S-A connectional axis, highlighting the predominant role of association connections in higher-order cognition^3,41–43^. Particularly, higher cognitive ability was associated with a more pronounced decline and lower connectivity strength in association connections compared to those with lower cognitive ability. This relationship may be mediated by the increased structural network segregation during development^3^. Together, these findings support spatially varying cognitive impacts on the maturation of SC across the human connectome, with the strongest effects in association connections.

Finally, we observed that the association between SC and psychopathology was also organized along the S-A connectional axis. Traditional diagnostic systems such as DSM-5^44^ rely on categorical classification that often fail to capture the spectrum characteristic of diseases and are marked by a high degree of comorbidity^45^. In contrast, dimensional frameworks such as the Research Domain Criteria^45,46^ emphasize transdiagnostic vulnerability, often captured by a general psychopathology factor (‘*p*-factor’)^29^. Our findings indicated that nearly all connections exhibited positive correlations between connectivity strength and *p*-factor scores, and the magnitude of these associations increased along the S-A axis. Participants with higher *p*-factor scores exhibited stronger connectivity strength for association connections. These results align with prior reports linking reduced segregation of higher-order association networks, such as the default mode and frontoparietal networks, to transdiagnostic psychopathology^47^. Moreover, higher *p*-factor levels were linked to earlier maturation and stronger connectivity strength of association connections. This pattern contrasted with the cognitive results, suggesting that atypical acceleration of association connections may represent a shared structural mechanism underlying both reduced cognitive performance and elevated psychopathology. From a translational perspective, the observed relationships between SC development and general psychopathology suggest a potential role of connectome-based normative frameworks in risk characterization. The developmental trajectories characterized here provide a reference against which individual developmental patterns of SC can be compared. Deviations from typical trajectories, particularly for connections localized at the association apex of the S-A connectional axis, may index neurodevelopmental vulnerability. The ConnectCharts platform could facilitate such individualized, trajectory-based assessment and may motivate further studies examining how deviations in SC strength relate to symptom change or treatment response^48^. Importantly, clinical translation will require prospective validation in clinically enriched cohorts and careful integration with behavioral and clinical assessments.

Several potential limitations should be noted. First, precisely reconstructing individual white matter connectivity is challenging, as dMRI tractography can produce false positives and negatives^49^. To mitigate these issues, we employed state-of-the-art probabilistic tractography with multi-shell, multi-tissue constrained spherical deconvolution^20^, together with ACT^21^ and SIFT^22^ to enhance biological accuracy. Consistency-based thresholding further reduced false-positive connections^50^. Previous studies have demonstrated the reliability of dMRI in tracing large-scale white matter bundles^51,52^, which was the focus of our system-level analyses. While such large-scale approaches are widely adopted in functional network studies^19,53–55^, they have rarely been applied to structural networks. Moreover, we confirmed the robustness of our findings by reconstructing connectomes from anatomically defined major bundles using TractSeg^25^. Second, recent advances in precision functional mapping highlight pronounced inter-individual variability in functional network boundaries, particularly within higher-order association cortex^53–57^. Future studies employing individual-level parcellations to define personalized SC may improve the interpretability of structure-function relationships by accounting for this topological variability.

Third, both the cognitive and psychopathological measures were composite measures, precluding inferences about domain-specific associations. Future investigations should untangle the relationships between SC and specific cognitive or psychopathological components. Fourth, although statistically significant, the effect sizes observed for both cognitive and psychopathological effects on SC were relatively small. Prior work has consistently demonstrated that effect sizes tend to be inflated in small samples, whereas larger samples provide more accurate estimates of the true effect size^58–60^. Nevertheless, the modest magnitude of the present SC-behavior associations limits their interpretability and precludes direct clinical translation. While the observed spatial organization of developmental effects across the connectome is robust at the group level, determining whether such patterns translate into clinically meaningful prediction or risk stratification will require prospective validation in clinically enriched and longitudinal samples. Additionally, the developmental variability patterns described here primarily reflects group-level effects. Although longitudinal data from the ABCD study were used, participants contributed at most two visits (baseline and two-year follow-up), limiting sensitivity to within-person developmental change and precluding detection of nonlinear transitions in the alignment between the spatial pattern of developmental rates and the S-A axis. Future studies with denser longitudinal sampling and longer follow-up periods may better capture within-person trajectories and their relationships with behavioral outcomes. Finally, our study primarily focused on understanding the developmental trajectories of SC between cortical systems. Future studies should extend this framework to subcortical and cerebellar circuits, which play crucial roles in cognitive and emotion^61,62^.

Notwithstanding these limitations, our findings provide compelling evidence that the maturation of SC follows a hierarchical program along the S-A connectional axis, linking macroscale structure to cognitive and psychopathological variability during youth. This developmental gradient underscores the importance of considering connectome-wide spatial variation in connectivity maturation when examining how brain maturation shapes behavior and vulnerability. By identifying connection-specific sensitive periods for plasticity, these results may inform the sensitive time windows for experiential, environmental, and interventional influences. To facilitate the exploration of these developmental patterns, we provided an interactive platform (http://connectcharts.cibr.ac.cn) enabling visualization of large-scale SC trajectories across the S-A connectional axis.

## Methods

### Participants

Our study utilized three independent neurodevelopmental datasets. The first one was a cross-sectional dataset from the Lifespan Human Connectome Project in Development (HCP-D)^17^. The HCP-D recruited typical developing participants aged 5 to 22 from four sites in the United States. We selected this dataset as the discovery dataset for developmental analyses, given its broad age range, high image quality, and consistent collection parameters across sites. Initially, demographic, cognitive, and neuroimaging data from 652 participants were obtained from the NIMH Data Archive (NDA) Lifespan HCP-D release 2.0. From this initial pool, we applied following exclusion criteria: 1) incomplete diffusion magnetic resonance imaging (dMRI) data; 2) anatomical anomaly; 3) under 8 years of age due to the small sample and big head motion often reported in this age group ^63^; 4) excessive head motion during dMRI scanning, identified by mean framewise displacement (FD) exceeding the mean plus three standard deviations (SD)^64^. Ultimately, we included 590 participants (273 males, aged 8.1–21.9) from the HCP-D. Written informed consent and assent were obtained from participants over 18 years of age and parents of participants under 18 years by the WU-Minn HCP Consortium. All research procedures were approved by the institutional review boards at Washington University.

The second dataset was from the Adolescent Brain Cognitive Development (ABCD) study^18^. The ABCD study recruited and followed approximately 10,000 children aged 9 to 10 years across the United States. Up to the beginning of data analyses of this study, the ABCD study had released neuroimaging data from the baseline and 2-year follow-up, covering ages 8.9 to 13.8 years. The sample is population based and includes participants with a range of psychiatric symptoms and diagnoses. We used ABCD as an independent replication dataset for developmental analyses and as the primary dataset for the cognitive and psychopathology analyses. We accessed neuroimaging data from the ABCD fast-tract portal in June 2022, and demographic, cognitive, and psychopathological measures from the ABCD release 5.1. The imaging data were acquired using scanners from SIEMENS, PHILIPS, or GE manufacturers. Our study exclusively utilized data from SIEMENS scanners, encompassing 5,803 scans from baseline and 4,547 scans from the 2-year follow-up, each including dMRI, associated field map, and T1-weighted imaging (T1WI). This decision aimed to mitigate bias from manufacturer variations and reduce computational costs. We selected the SIEMENS manufacturer because most data were collected by scanners from this manufacturer in the ABCD study. From these scans, we applied various exclusion criteria including: 1) not meeting the official imaging recommended inclusion criteria outlined in the release 4.0 notes (we adopted criteria from release 4.0 because release 5.1 was not available when the MRI processing was conducted.); 2) incomplete dMRI data or failure in unzip or format conversion process; 3) lack of parental fluency in English or Spanish; 4) lack of proficiency in English; 5) diagnosis of severe sensory, intellectual, medical or neurological issues; 6) prematurity or low birth weight (N = 2,350); 7) having contraindications to MRI scanning; 8) invalid data regarding age and sex; 9) failure in data processing; 10) excessive head motion (mean FD > Mean + 3SD). The criteria regarding demography and healthy conditions came from a prior study^65^. After applying these criteria, we included a total of 7,104 eligible scans for the subsequent analyses, comprising 3,949 from baseline (2,075 males, aged 8.9–11.0) and 3,155 from 2-year follow-up (1,701 males, aged 10.6–13.8). The study protocol was approved by the institutional review board of the University of California, San Diego. Before participation, parents or legal guardians provided written informed consent, and children provided verbal assent.

The third dataset comprised three Chinese studies, collectively referred to as the Chinese Cohort, used to test generalizability in an Asian sample. The Executive Function and Neurodevelopment in Youth (EFNY) study is an ongoing project recruiting typically developing youth and youth with attention or learning disorders in Beijing. Here, we used EFNY data collected through December 10, 2025. The second study, the developmental component of the Chinese Color Nest Project (devCCNP)^66,67^ focused on longitudinal developmental research with typically developing youth. Only the Beijing site of the devCCNP was included because dMRI data were available. The third part was the healthy control group of the Shandong Adolescent Neuroimaging Project on Depression (SAND)^68,69^, which recruited youth from Shandong Province, China. From the three studies, we initially obtained 1,082 scans with complete dMRI and T1WI data (EFNY: N = 547; devCCNP: N = 384; SAND: N = 151). Then, we applied the following exclusion criteria: 1) severe sensory, intellectual disability (IQ under 70), medical or neurological issues; 2) prematurity or low birth weight; 3) missing or invalid age or sex information; 4) data processing failures; 5) under 6 years of age due to the small sample size; 6) excessive head motion (mean FD > Mean + 3SD). Finally, we included 947 scans (481 males, aged 6.1–23.4) for the subsequent analyses, in which 490 scans from the EFNY, 312 scans from the devCCNP (59 participants have multiple measurements), and 145 scans from the SAND. Written assent and informed consent were obtained from all participants and, for those under 18 years old, from their parents or legal guardians. Ethical approvals were obtained as follows: the devCCNP was approved by the Institutional Review Board of the Institute of Psychology, Chinese Academy of Sciences; the EFNY by the Human Research Ethics Committee of the Chinese Institute for Brain Research, Beijing; and the SAND by the Institutional Review Boards at Shandong Normal University.

The HCP-D, ABCD, and Chinese Cohort participant inclusion and exclusion flow charts are presented in **Fig. S1**–S**3**. Additional demographic details for the included datasets are available in **Table S1**–**S3**.

### Cognitive assessment

The HCP-D and ABCD studies assessed participants’ cognitive abilities using NIH Toolbox Cognition Battery. This battery evaluates five fluid cognitive functions: flanker inhibitory control and attention, list sorting working memory, dimensional change card sort, picture sequence memory, and pattern comparison processing speed. For the HCP-D and ABCD datasets, a fluid composite score reflecting participants’ higher-order cognition was computed from these five fluid cognitive measurements^38^. Specifically, based on the NIH Toolbox national norms, raw scores from each task were converted into normally distributed standard scores, with a mean of 100 and a standard deviation of 15. These standardized scores were then averaged, and the resulting average score was re-standardized to acquire the fluid composite score^26^. Notably, uncorrected standard scores (without age adjustment) were used in the primary analyses, and age-corrected standard scores were used in a sensitivity analysis to confirm that our findings were not dependent on the score normalization approach.

### Psychopathology assessment

Prior research identified a general psychopathology factor (also referred to as ‘*p*-factor’), which represents a shared vulnerability to broad psychiatric symptoms and accounts for the comorbidity across mental disorders^29^. The ABCD dataset provides a large, transdiagnostic, population-based sample spanning healthy youth, those at elevated risk, and youth with at least one diagnosis, making it well suited for our psychopathology analyses. We indexed symptoms using parent-report Child Behavior Checklist (CBCL)^70^ scores. The CBCL contains 113 items covering emotional and behavioral domains aligned with DSM classifications and has strong psychometric properties, supporting its extensive use in clinical and research settings^71^.

Building on Moore et al.^27^, who specified a CBCL bifactor structure in ABCD, we estimated a confirmatory bifactor model in the full ABCD Release 5.1 sample using Mplus 8.3^72^ (baseline N = 11,860; 1-year N = 11,201; 2-year N = 10,895; 3-year N = 10,095; 4-year N = 4,679). Following that structure, 66 CBCL items were modeled to load on four orthogonal dimensions: a general psychopathology (“*p*-”) factor plus three specific factors—internalizing, attention-deficit/hyperactivity disorder (ADHD), and conduct problems (**Fig. 7e**). To minimize bias from selective participation or visit timing on item loadings, we fit the model to the entire sample, stratified by site, accounted for clustering within families, and constrained factor loadings to be equal across time points (metric invariance). Model fit met conventional benchmarks^73^: comparative fit index (CFI) = 0.96 (> 0.90), root mean square error of approximation (RMSEA) = 0.02 (< 0.08), and standardized root mean square residual (SRMR) = 0.07 (< 0.08). The resulting *p*-factor scores provide a continuous, transdiagnostic index of symptom burden across development. To evaluate robustness to psychopathology operationalization, we additionally used the CBCL Total Problems raw score as an alternative measure of overall psychiatric symptom burden. The Total Problems score was calculated by summing parent-reported symptom ratings across all CBCL problem items, with higher scores indicating greater overall symptom burden.

### MRI acquisition

T1WI and dMRI data were acquired for each participant in this study. MRI data for the HCP-D, ABCD, Chinese Cohort-EFNY, and Chinese Cohort-SAND datasets were obtained using 3T SIEMENS scanners. For the Chinese Cohort-devCCNP, MRI data were acquired using a GE Discovery MR750 3T scanner. Detailed imaging acquisition parameters are summarized in **Table S4**.

Minimally processed T1WI data were obtained from the HCP-D dataset, while SIEMENS-normalized T1WI data were obtained from the ABCD dataset and the EFNY study in Chinese Cohort. Raw T1WI data were obtained from the other studies within the Chinese Cohort. The minimally processed T1WI data from the HCP-D dataset underwent gradient distortion, anterior commissure-posterior commissure (ACPC) alignment, and readout distortion correction^74^. For dMRI, raw data were obtained from all datasets.

### MRI data processing

Initially, we applied the anatomical pipeline embedded in QSIPrep version 0.16.0 (https://qsiprep.readthedocs.io/)^75^ to the T1WI data from all the datasets. QSIPrep is an integrative platform for preprocessing dMRI and structural imaging data, as well as reconstructing white matter structural connectome^75^ by incorporating tools from FSL^76^, DSI Studio (https://dsi-studio.labsolver.org/), DIPY^77^, ANTs (https://stnava.github.io/ANTs/), and MRtrix3^78^. The anatomical pipeline conducted through ANTs included: 1) intensity non-uniformity correction; 2) removal of non-brain tissues; 3) normalization to the standard Montreal Neurological Institute (MNI) space. The skull-stripped T1WI in native space was used as the anatomical reference for the dMRI workflow. Normalization generated transformation matrices to register the atlas in MNI space to individual anatomical references.

Next, we utilized the intensity normalized T1WI data from all the datasets to reconstruct surface and segment tissues through *FreeSurfer*^79^ (http://surfer.nmr.mgh.harvard.edu/). The surface pial and tissue segmentations would be used as anatomical constraints during the construction of the structural connectome. The HCP-D T1WI was processed based on the *FreeSurfer* workflow from the HCP processing pipelines^74^, while the T1WI from the other datasets were processed using the recon-all pipeline through *FreeSurfer* version 7.1.1.

We applied the dMRI pipeline embedded in QSIPrep to the dMRI data from all the datasets. The pipeline included: 1) aligning and concatenating runs of dMRI and associated field maps; 2) designating frames with a b-value less than 100 s/mm^2^ as b = 0 volumes; 3) Marchenko-Pastur principal component analysis (MP-PCA) denoising through MRtrix3’s *dwidenoise* function^80^; 4) Gibbs unringing through MRtrix3’s *mrdegibbs* function^81^; 5) B1 bias correction through MRtrix3’s *dwibiascorrect* function^82^; 6) head motion, distortion and eddy current corrections through FSL’s eddy tool^83^; 7) coregistration to individual T1WI and realignment to ACPC orientation. During this process, distortion correction utilized b = 0 reference images with reversed phase encoding directions. Since the devCCNP and SAND studies did not scan the field map for dMRI, distortion correction was not applied to the data from these two studies. Following the previous study^84^, the mean FD was calculated as the sum of the absolute values of the differentiated realignment estimates for each volume to estimate the head motion during the dMRI scans.

### Reconstruction of structural connectome

For each scan, whole-brain probabilistic fiber tracking was performed using MRtrix3^78^. In the HCP-D, ABCD, and Chinese Cohort-EFNY datasets, dMRI were scanned using multi-shell parameters, so multi-shell multi-tissue constrained spherical deconvolution (MSMT-CSD)^20^ was utilized to estimate the fiber orientation distribution (FODs) for each voxel. In devCCNP and SAND, dMRI data were scanned using single-shell parameters, so we used single-shell 3-tissue CSD (SS3T-CSD)^85^ to estimate FODs. To improve biological plausibility, we applied anatomically constrained tractography (ACT) with hybrid surface–volume segmentation (HSVS)^21,86^, leveraging FreeSurfer-derived pial surfaces and tissue segmentations. FOD-guided probabilistic tracking (iFOD2) generated 10 million streamlines per scan, restricted to lengths of 30–250 mm via *tckgen* function^87^. Finally, to align streamline densities with fiber densities implied by the FODs, we assigned per-streamline weights using SIFT2 via *tcksift2* function^22^.

Large-scale SC connectomes with larger nodes have demonstrated higher reproducibility and biological validity compared to those with finer nodes^88–90^. Therefore, we parcellated the cortex into 12 large-scale systems along the sensorimotor-association (S-A) cortical axis, which was derived from multiple cortical features to capture a cortical hierarchical gradient from lower-order unimodal areas to higher-order transmodal areas^10^. Specifically, we first ordered the brain regions from the Schaefer-400 atlas^91^ along the S-A cortical axis and removed limbic regions due to their low signal-to-noise ratio^92^, yielding 376 regions. These regions were then grouped into 12 large-scale systems, with each system comprising 31 or 32 regions. The number 12 was chosen as the median value between the 7- and 17-system parcellations commonly used in previous large-scale brain network studies^19,93^. Additionally, we included 7- and 17-system configurations in sensitivity analyses to assess the robustness of our findings. A radial search (maximum radius = 2 mm) was performed from each streamline endpoint to identify the nearest gray matter node. To reduce false-positive streamlines in global tractography^49^, we applied a consistency-based thresholding method^94^ on the region-level connectome. Streamlines belonging to connections with a coefficient of variation (CV) above the 75th percentile were excluded^50,95^. Finally, for each participant at each visit, we constructed a 12×12 large-scale network with 78 undirected connections. The weight of each connection, defined as SC strength, was calculated by multiplying the number of streamlines by the SIFT2 coefficients, and then dividing by the arithmetic mean volume of the connected system pair.

### Statistical analysis

#### Development of structural connectivity strength in youth

We first evaluated the connectome-wide spatial variation in developmental trajectories of SC strength during youth. All statistical analyses were performed in R4.1.0. We fitted the developmental models for 78 large-scale structural connections in all the three datasets. We utilized generalized additive models (GAMs) for the cross-sectional datasets (HCP-D and Chinese Cohort) and generalized additive mixed models (GAMMs) for the longitudinal ABCD dataset to flexibly capture linear and non-linear age-related changes in SC strength. Notably, although 59 participants from the devCCNP study in Chinese Cohort have more than one measurement, their proportion (6.8%) is so small that we treated the Chinese Cohort dataset as a cross-sectional dataset. The models were fitted using *mgcv* ^96^ and *gamm4* package^97^. For each model, we set SC strength as the dependent variable, with age as a smooth term, and sex and mean FD as covariates, as shown in equation (1). Per-participant random intercepts were additionally included in the GAMMs. Smooth plate regression splines served as the basic function of the smooth term, and the restricted maximal likelihood approach was used to estimate smoothing parameters. To determine the optimal degree of freedom (*k*) for the smooth term in the GAM/GAMM models, we evaluated model fit using the Akaike Information Criterion (AIC) across *k* values ranging from 3 to 6. We conducted a stratified bootstrap analysis with 1,000 iterations, resampling participants with replacement while preserving site proportions. In each iteration, we compared AIC values across *k* values for 78 structural connections and identified the optimal *k* as the value most frequently selected across connections. A *k* value of 3 emerged as optimal in 773 iterations for the HCP-D dataset, all 1,000 iterations for ABCD, and 617 iterations for the Chinese Cohort (**Fig. S14**). Therefore, we determined the *k* value as 3, which is also consistent with previous neurodevelopmental studies in youth^11,12^.

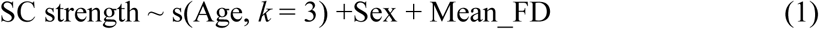

For each model, we evaluated the significance of the age effect by comparing the full model with a nested model without age^98^, using parametric-bootstrap ANOVA for GAMs (via *ecostats*^99^) and parametric-bootstrap likelihood-ratio tests for GAMMs (via *pbkrtest*^100^), each with 1,000 simulations. The *P* values were then adjusted using the false discovery rate (FDR) correction, with a significant threshold set at 0.05. We calculated the first derivative of the age smooth function for each model using the *gratia* package^101^ to assess the change rate of SC strength. The first derivatives were computed for 1,000 age points sampled at equal intervals within the age span. For derivatives at each age point, we computed the *P* values of the derivatives based on the 95% CI and then adjusted the *P* values using the FDR method for 78 models. Age windows of significant development were identified when the first derivative had a *P_FDR_* < 0.05. To assess the overall age effect, we calculated the partial *R*^2^ between the full and null models and then assigned the sign based on the average first derivative of the smooth function^12^. We further examined within-person developmental effects using ABCD participants with both baseline and two-year follow-up data (N = 2,570). We decomposed age into a between-person component (age_bp, baseline age) and a within-person component (age_wp, the within-person age interval relative to baseline). We then estimated within-person age effects using linear mixed-effects models (equation 2). The *T* values for the age_wp coefficient were extracted as connection-wise estimates of within-person age effects.

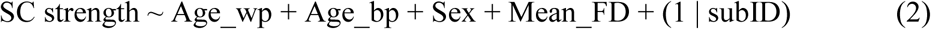

To visualize the developmental trajectory of SC strength for each connection, we predicted the model fits for 1,000 age points sampled at equal intervals within the age span. Covariates were set at the median for numerical variables and the mode for categorical variables. The SC strength was normalized by dividing the initial strength at the youngest age (HCP-D: 8.1 years; ABCD: 8.9 years; Chinese Cohort: 6.1 years) within the age span for visualization, termed as SC strength ratio. As shown in **Fig. 2b**, we observed heterogeneity in the curvature of the developmental trajectories. To highlight this heterogeneity, we standardized the fits (z-scored) across the age span for each connection.

As we observed heterogeneity in the curvature of developmental trajectories, we computed the average second derivatives of age via central finite differences to quantify this curvature. A positive value of the average second derivative indicates a concave upward trajectory, while a negative value indicates a concave downward trajectory. The greater the absolute value, the greater the degree of curvature. Notably, there is significant variability in the average weights of SC strength across the connectome, ranging from 0.4 to 12.3. It is important to recognize that a higher first or second derivative does not necessarily imply a greater change rate or curvature of trajectory relative to the connection’s initial strength. For instance, a connection starting with an initial strength of 0.5 and a first derivative of 0.1 will evolve faster than a connection with an initial strength of 5 and a first derivative of 0.2, relative to their starting points. To address this issue, we normalized the SC strength of each connection by its fitted value at the youngest age within the dataset’s age span, resulting in a SC strength ratio relative to its initial strength. We then computed the first and second derivatives of the models with the SC strength ratio as the dependent variable. This normalization process does not alter the significance or magnitude of the age effect.

To compare developmental statistics across connection types, we classified the 12 systems into sensorimotor (“S”) and association (“A”) groups based on their overlap with the Yeo-7 networks: the four systems closest to the sensorimotor pole (most overlapping with visual and somatomotor networks) were assigned to S, and the remaining eight systems to A. The 78 connections were then categorized as SS, SA, or AA. We used Cohen’s d to quantify (i) differences in age effects between pairs of connection types and (ii) whether the second derivatives within each connection type differed from zero. Statistical significance was assessed using permutation testing by shuffling connection-type labels 1,000 times to generate a null distribution of Cohen’s d. Permutation *P* values were defined as the proportion of permuted Cohen’s d values at least as extreme as the observed value.

#### Definition of the S-A connectional axis

The S-A cortical axis provided a framework to characterize the heterochrony of postnatal regional neurodevelopment, suggesting that many cortical features progress along the S-A cortical axis during childhood and adolescence^11–13^. Building upon this, we aimed to investigate whether the development of SC strength unfolds along the S-A axis in terms of connections. As shown in equation (3), we defined the element (*C_i,j_*) between *node_i_* and *node_j_* in the matrix of the S-A connectional axis as the quadratic sum of the S-A cortical axis ranks of *node_i_* and *node_j_*. Next, we computed the ranks of *C* as the S-A connectional axis rank.

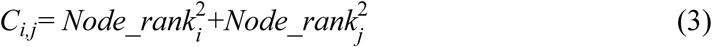

In the context of a 12×12 large-scale connectome, the S-A connectional axis rank ranges from 1 to 78. To address potential concerns about arbitrariness in defining the S-A connectional axis, an alternative definition of the S-A connectional axis by multiplying the S-A cortical axis ranks of each pair of nodes was validated (**Fig. S10c**; **Supplementary text**).

#### Developmental alignment with the S-A connectional axis

The main goal of this study is to determine if the spatial variation of SC development aligns with the S-A connectional axis. To achieve this, we utilized Spearman’s rank correlation to evaluate the concordance and its significance between the S-A connectional axis rank and both 1) the magnitude and direction of developmental effects (partial *R*^2^) and 2) the curvature of the developmental trajectories (average second derivative). To depict developmental trajectories along the S-A connectional axis continuously, we divided the S-A connectional axis into 10 decile bins, with each bin consisting of 7 or 8 large-scale structural connections. We then calculated the average trajectory within each bin and subsequently normalized these averages using z-scores (**Fig. 3d**).

To comprehensively understand how alignment evolves across the youth, we performed an age-resolved analysis of the alignment between the developmental change rates of connectivity strength and the S-A connectional axis^12,13^. We calculated the first derivative to measure the developmental change rates. This approach enabled us to capture the evolving alignment of development with the S-A connectional axis across the targeted age span. To determine the correlation coefficient and 95% CI for these age-specific correlation values, we initially sampled 1,000 times from the specified multivariate normal distribution of the independent variables’ coefficients for each connectional model. We then generated the posterior derivatives at 1,000 age points based on the posterior distribution of each connectional fitted model. Subsequently, we repeated the process of correlating the S-A connectional axis rank with 1,000 draws of the posterior derivative of the age smooth function at each of the 1,000 age points. The resultant distribution of correlation coefficients was utilized to determine the median and 95% CI of alignment at each age point. Additionally, we employed the sampling distribution of age-specific S-A connectional axis correlation values to identify the age at which the alignment flipped from negative to positive. This involved calculating the age at which the axis correlation was closest to zero across all 1,000 draws and reporting the median along with the 95% CI.

To examine potential sex differences in the timing of the developmental shift in the alignment between spatial patterns of SC developmental rates and the S-A connectional axis, we conducted sex-stratified analyses in the HCP-D dataset. GAMs were fitted separately in females (N = 317) and males (N = 273) for each of the 78 connections. Using the fitted models, we estimated the age at which the alignment between SC developmental rates and S-A axis crossed zero, following the procedure described above. We then computed the observed sex difference as the difference in the median zero-crossing age between females and males. To assess the statistical significance of this difference, we performed a participant-level permutation test. Sex labels were randomly shuffled across participants 1,000 times, and for each permutation the full sex-stratified modeling and alignment procedure was repeated to generate a null distribution of differences in median zero-crossing age. The observed sex difference in median zero-crossing age was compared with this null distribution to obtain a permutation-based *P* value.

Based on the age-resolved analysis, we found a transition age of 15.5 years at which the alignment of developmental effects with the S-A connectional axis shifts from negative to positive in the HCP-D dataset. To test whether the overall SC developmental effects differ spatially before and after this transitional age, we split all participants into two subsets (younger subset: *N* = 355, aged 8.1–15.5 years; older subset: *N* = 235, aged 15.5–21.9 years). We re-evaluated the developmental effects (partial *R*^2^) of SC strength for the two subsets separately using GAMs with the same parameters as those used for the full sample. Then, we utilized Spearman’s rank correlation analysis to test the alignment of the overall developmental effects (partial *R*^2^) with the S-A connectional axis in the two subsets. Notably, we expected the replicated results from the ABCD dataset to be consistent with those found in the younger subset of the HCP-D because the age span of ABCD participants (8.9–13.8 years) was within the 8.1 to 15.5 years range.

#### Associations between structural connectivity strength and higher-order cognition

Associations between the fluid cognition composite score and SC strength were examined within the HCP-D and ABCD datasets. Cognitive analyses were not applied to the Chinese Cohort dataset because the three studies within this cohort utilized different cognitive tasks. Additionally, due to the lack of measurements for working memory and flexibility, which are components of the cognitive score, in the 2-year follow-up data of the ABCD study, only baseline data from the ABCD dataset were included in the cognitive analyses. We employed GAMs to assess the relationships between the SC strength and the fluid composite score for each connection, controlling for age, sex, and mean FD. The equation for GAMs is shown below (equation (4)). For the sensitivity analysis using age-corrected cognitive scores, the smooth term of age was removed to avoid double adjustment^102^.

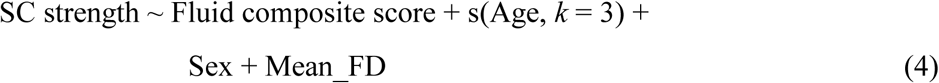

The *T* values of the fluid composite score indicate the magnitude and direction of the association, and its significance was determined by comparing the full model with a null model lacking the cognition term. GAM comparisons utilized parametric bootstrap testing via analysis of variance with 1,000 simulations, facilitated by the *ecostats* package^99^. The *P* values were then FDR corrected across all the 78 connections. We next evaluated the alignment of association magnitude and direction with the S-A connectional axis rank across all connections through Spearman’s correlation analysis.

Furthermore, we depicted the developmental trajectories by different levels of the cognitive composite score to elucidate how SC strength evolved in individuals with varying cognition levels. To do this, we fitted an age-by-cognition interaction model for each connection controlling for sex, and mean FD (see equation (5) for the formula).

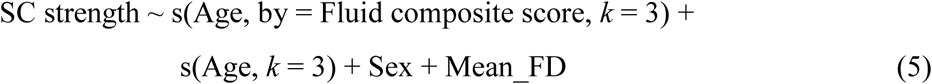

For this age-by-cognition interaction analysis, we included both the observations from baseline and 2-year follow-up of the ABCD dataset and utilized baseline cognitive score as the by-item to fit GAMM models. Using the acquired models, we estimated SC strength by assigning cognitive scores as low and high levels respectively. To define these levels, we used the 10th percentile of average cognitive performance for the low level, and the 90th percentile for the high level. We then averaged trajectories for low and high cognition levels independently within deciles of the S-A connectional axis for visualization purposes.

#### Psychopathological associations with structural connectivity strength

We further evaluated the associations between the general psychopathological factor, *p*-factor, and SC strength. The associations were evaluated using the ABCD dataset through GAMMs while controlling age, sex, mean FD and per-participant random intercepts. See below for the equation (6).

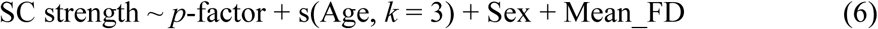

Similar to cognitive analysis, the *T* value of the *p*-factor term indicates the magnitude and direction of the association between *p*-factor and SC, and its significance was determined by comparing the full model with a null model lacking the *p*-factor term. GAMM comparisons employed a parametric bootstrap method utilizing the likelihood ratio test statistic with 1,000 simulations, supported by the *pbkrtest* package^100^. FDR correction was utilized to adjust *P* values. To evaluate the alignment of psychopathological associations with the S-A connectional axis rank, we performed Spearman’s rank correlation analysis.

Subsequently, we further depicted the developmental trajectories by different levels of *p*-factors to elucidate how developmental trajectories of SC strength differed between individuals with varying severity of general psychiatric symptoms. To achieve this, we modeled age-dependent changes in SC strength as a function of *p*-factors (see equation (7)).

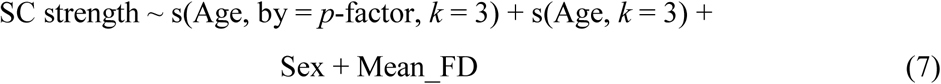

Based on this model, we estimated SC strength by assigning *p*-factor as low and high levels respectively. To define these levels, we used the 10th percentile of the *p*-factor for the low level, which represents no or mild psychiatric symptoms, and the 90th percentile for the high level, which represents severe psychiatric symptoms. We then averaged trajectories for low and high *p*-factor levels independently within deciles of the S-A connectional axis for visualization purposes.

#### Correction for multi-site batch effects

Data from the three datasets were collected across multi-acquisition sites with varied scanners and sequences. To harmonize SC strength across acquisition sites, we applied ComBat-GAM^103^ and NonlinearLongitudinalComBat (https://github.com/nmd1994/NonlinearLongitudinalComBat) to the 78 SC connections derived from diffusion MRI data. ComBat-GAM was used for the cross-sectional datasets, whereas NonlinearLongitudinalComBat was applied to the longitudinal ABCD dataset. Both approaches effectively remove site-related batch effects while preserving biological phenotypic effects, particularly nonlinear age-related effects, making them well suited for developmental datasets. During the harmonization process, covariates included age, sex, and head motion, with age specified as a smooth term. For analyses involving cognitive and psychopathological measures, the corresponding cognitive and psychopathological variables were additionally included as covariates to ensure that these phenotypic effects were preserved during harmonization, following established practice in prior studies^104–106^.

#### Sensitivity analyses

We performed a series of sensitivity analyses to ascertain the robustness of our findings. The detailed method description was presented in the **Supplementary text**.

## Data availability

The HCP-Development 2.0 Release data used in this report came from DOI: 10.15154/1520708 via the NDA (https://nda.nih.gov/ccf). The ABCD 5.1 data release used in this report came from DOI: 10.15154/z563-zd24 via the NDA (https://nda.nih.gov/abcd). The fast-track data from the ABCD Study data is also available through the NDA. The devCCNP data in the Chinese Cohort is available via the Science Data Bank (https://doi.org/10.57760/sciencedb.07478). However, data from the EFNY and SAND studies are available under restricted access because data collection for both datasets is still ongoing. The principal investigator of EFNY plans to release the raw data after the collection of Phase I is completed.

## Code availability

All codes used to perform the analyses in this study and the statistical magnitudes derived from analyses can be found at https://github.com/CuiLabCIBR/SCDevelopment.git. To enhance accessibility, we also developed an interactive website (https://connectcharts.cibr.ac.cn/) that allows the broader community to explore and use our developmental charts. All analysis methods are described in the main text and supplementary materials.

## Acknowledgments

We thank the research participants and staff involved in data collection and project executive of the Lifespan Human Connectome Project in Development (HCP-D) study, Adolescent Brain Cognitive Development (ABCD) Study, the Executive Function and Neurodevelopment in Youth (EFNY), the developmental component of the Chinese Color Nest Project (devCCNP), and the Shandong Adolescent Neuroimaging Project on Depression (SAND). The HCP-D data were provided by the Human Connectome Project, WU-Minn Consortium (Principal Investigators: David Van Essen and Kamil Ugurbil; 1U54MH091657) funded by the 16 National Institutes of Health (NIH) Institutes and Centers that support the NIH Blueprint for Neuroscience Research; and by the McDonnell Center for Systems Neuroscience at Washington University. Research reported in this publication was supported by the National Institute of Mental Health of NIH under Award Number U01MH109589 and by funds provided by the McDonnell Center for Systems Neuroscience at Washington University in St. Louis. The ABCD data were provided by the ABCD Study (abcdstudy.org) funded by NIH. The ABCD Study is a multisite, longitudinal study designed to recruit more than 10,000 children ages 9-10 years old and follow them over 10 years into early adulthood. The ABCD Study is supported by the NID and additional federal partners under award numbers U01DA041048, U01DA050989, U01DA051016, U01DA041022, U01DA051018, U01DA051037, U01DA050987, U01DA041174, U01DA041106, U01DA041117, U01DA041028, U01DA041134, U01DA050988, U01DA051039, U01DA041156, U01DA041025, U01DA041120, U01DA051038, U01DA041148, U01DA041093, U01DA041089, U24DA041123 and U24DA041147. A full list of supporters is available at https://abcdstudy.org/federal-partners.html. A listing of participating sites and a complete listing of the study investigators can be found at https://abcdstudy.org/consortium_members/. ABCD consortium investigators designed and implemented the study and/or provided data but did not necessarily participate in the analysis or writing of this report. This manuscript reflects the views of the authors and may not reflect the opinions or views of the NIH or ABCD consortium investigators. The ABCD data repository grows and changes over time. The Executive Function and Neurodevelopment in Youth data were provided by our lab in CIBR, supported by the Brain Science and Brain-like Intelligence Technology - National Science and Technology Major Project (Grant No.2022ZD0211300). The CCNP data were contributed by the CCNP consortium and funded by the Start-up Funds for Leading Talents at Beijing Normal University, the Key-Area Research and Development Program of Guangdong Province (Grant No.2019B030335001), and the National Basic Science Data Center “Chinese Data-sharing Warehouse for In-vivo Imaging Brain” (Grant No.NBSDC-DB-15). The SAND data were provided by K.W. from the School of Psychology at Shandong Normal University and supported by the National Natural Science Foundation of China (Grant No.32000760).

## Author contributions

Z.C. and X.X. conceptualized the study. X.X., H.Y., and J.C. curated the data. X.X., J.C., H.X., J.K., S.Z., T.X. analyzed the data. H.S., T.X., and F.Y. provided guidance on data analysis and interpretation of the results. Z.C. managed the project administration and supervised the project. X.X. and Z.C. wrote the original draft. V.J.S. and F.Y. commented on the manuscript. X.X., Z.C. and H.X. designed the interactive platform. Z.C., Y.L. and K.W. provided part of data. J.K. and T.X. facilitated the sensitivity analyses. Z.C., X.X., H.Y., J.C., and V.J.S. reviewed and edited the manuscript.

## Supplemental information

Supplementary text

Fig. S1 to S14

Tables S1 to S4

