## Supplementary Materials for "Mapping the spatiotemporal continuum of structural connectivity development across the human connectome in youth"

**Supplemental information**

Supplementary text
Fig. S1 to S14
Tables S1 to S4

### Supplementary text

#### Methods of sensitivity analyses

We conducted sensitivity analyses to show that the developmental variability was robust to methodological choices, including 1) varying the number of sensorimotor-association (S-A) cortical systems; 2) reconstructing structural connectomes using the canonical Yeo-7 and Yeo-17 cortical parcellations; 3) regressing out Euclidean distance between system pairs when assessing S-A alignment; 4) controlling for mean whole-brain SC strength; 5) including socioeconomic status (SES) and intracranial volume (ICV) as additional covariates; 6) rebuilding connectomes from major bundle-based TractSeg tractography; and 7) defining an alternative S-A connectional axis based on the product of system ranks. In addition, we tested whether the S-A connectional axis captures the dominant developmental organization when the developmental axis is defined in a data-driven manner, and compared its explanatory power with that of other organizational principles. The developmental analyses were replicated using the HCP-D dataset. For each sensitivity analysis, we examined the alignments of second derivatives of developmental trajectories and age-resolved developmental slopes with the S-A connectional axis.

The first sensitivity analysis examined whether the scale of the large-scale structural connectome affected the results. Rather than parcellating the cortex into 12 cortical systems, we generated cortical parcellations with 7 or 17 systems defined along the S-A axis, ensuring each system contained approximately the same number of brain regions. We then reconstructed structural connectomes with 28 connections among 7 systems and 153 connections among 17 systems for each scan. Developmental models were refit for each connection in the HCP-D dataset.

Second, to assess robustness using canonical Yeo functional system parcellations[1], we constructed large-scale structural connectomes based on the Yeo-7 or Yeo-17 systems defined from the Schaefer-400 atlas. Consistent with our primary analysis, limbic regions were excluded, leaving 6 systems in the Yeo-7 parcellation and 15 systems in the Yeo-17 parcellation. We ranked the Yeo systems according to the mean S-A cortical-axis rank of the regions within each system. The Yeo-7 connectome comprised 21 undirected connections, whereas the Yeo-17 connectome comprised 120 undirected connections. We then refit the developmental models for each connection in the HCP-D dataset.

Third, because Euclidean distance between regions can influence the developmental patterns of structural connections, we evaluated whether this distance might confound our findings. Specifically, we regressed the Euclidean distance between each pair of systems in MNI space out of the average second derivatives of developmental trajectories and age-resolved developmental slopes. We then repeated the analyses examining the alignment of these metrics with the S-A connectional axis.

The fourth sensitivity analysis assessed whether the observed developmental variations in SC persisted after controlling for global connectivity strength. To this end, we computed mean whole-brain SC strength by averaging connection strengths across all 78 system-to-system connections in the 12 × 12 large-scale connectome. Mean whole-brain SC strength was then included as an additional covariate when fitting the developmental models for each individual connection.

Fifth, we tested whether our findings were confounded by SES or ICV. SES was measured using the family income-to-needs ratio, as in prior work[2,3]. The family income-to-needs ratio was calculated as annual family income divided by the federal poverty threshold, as defined by the U.S. Department of Health and Human Services (https://aspe.hhs.gov/topics/povertyeconomic-mobility/poverty-guidelines/prior-hhs-poverty-guidelines-federal-register-references). Intracranial volume was computed via *FreeSurfer*. We refitted the developmental models for each connection while additionally controlling for the participants' SES or ICV.

Sixth, we reconstructed the "global tractography" by merging 72 anatomically well-described bundles from TractSeg for each participant, which helped to minimize potential noise introduced by global tractography[4]. TractSeg is a convolutional neural network-based approach that accurately and rapidly segments tracts in individual spaces[5]. It segments white matter bundles and their end regions, which are then used for bundle-specific tractography. The structural connectomes reconstructed based on the major bundle-based tractography contain only long fibers belonging to the major tracts such as arcuate fascicle and corpus callosum. Specifically, we extracted the peaks of the constrained spherical deconvolution (CSD) function of white matter, identifying bundle start and end segmentations for all 72 bundles using TractSeg. These bundles were reconstructed using the iFOD2 algorithm, with an upper limit of 10k streamlines generated for each bundle. Consistent with the primary analysis, we constructed structural connectomes based on 12 systems along the S-A cortical axis. Structural connectivity strength was estimated by multiplying streamline counts by SIFT2 coefficients, normalized by the average volume of the paired systems. Streamlines belonging to connections with a coefficient of variation (CV) above the 75th percentile were excluded. Although this method excludes short regional u-fibers, which are biologically feasible, it significantly reduces potential noise from spurious fibers. We re-estimated the developmental models for connections derived from the major bundle-based tractography to test whether the observed developmental variability along the S-A axis was influenced by tractography noise.

Seventh, in the primary analyses, we defined the S-A connectional-axis rank of each edge as the squared sum of the S-A cortical-axis ranks of the connected systems. To demonstrate that developmental patterns of structural connections are consistent across alternative definitions of the S-A connectional axis, we instead defined this rank as the product of the S-A cortical-axis ranks of each pair of systems (**Fig. S10c**). Under this alternative definition, connections between intermediate-level areas are assigned higher ranks than connections between the most sensorimotor and most transmodal association regions. We then repeated the developmental analyses using this alternative definition.

Finally, to derive a data-driven dominant axis of SC developmental variation, we applied principal component analysis (PCA) to connection-wise trajectories of SC developmental rates (**Fig. 4a**; **Fig. S11a**). Specifically, we assembled the change-rate trajectories into a matrix with connections as rows and age points as columns, and performed PCA on this matrix. The loadings of the first principal component (PC1) across connections were taken as the dominant, data-driven developmental axis. We then assessed the spatial correspondence between this developmental axis and the S-A connectional axis, as well as alternative connectional axes derived from principal functional gradient and T1w/T2w measures, using Spearman's rank correlation.

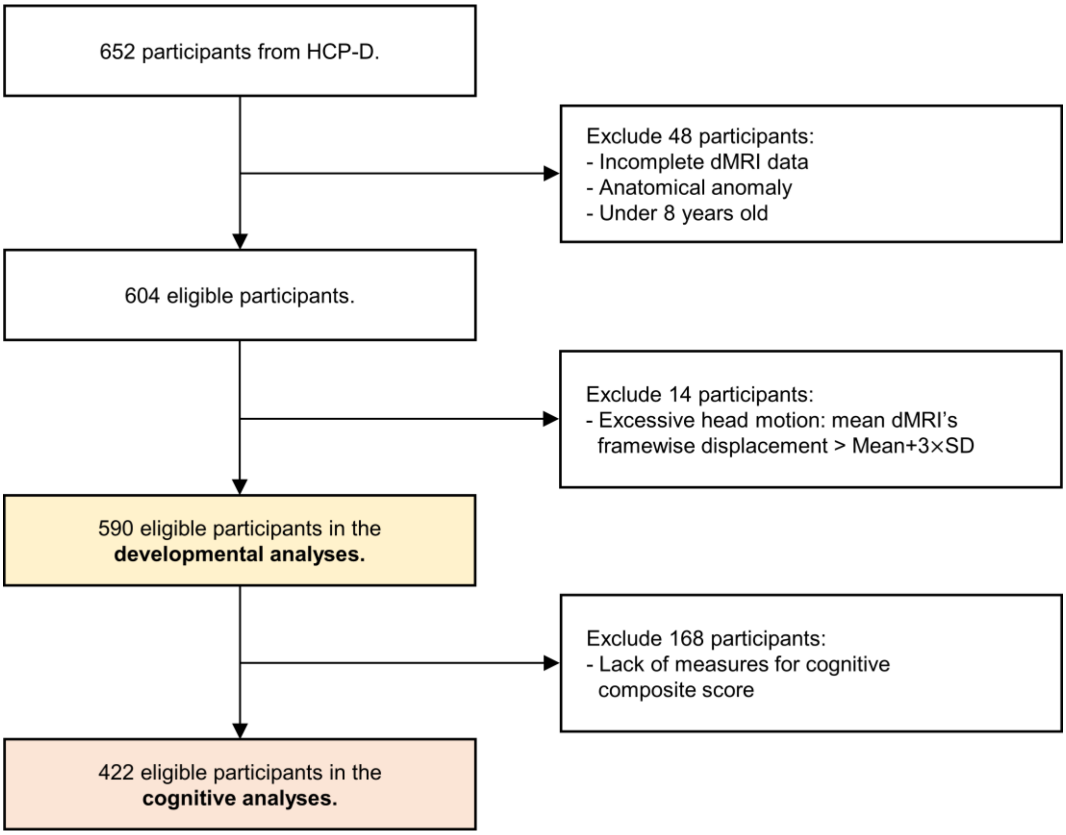


**Fig. S1. Flowchart of inclusion and exclusion for participants in the HCP-D dataset.** HCP-D: the Lifespan Human Connectome Project in Development; dMRI: diffusion magnetic resonance imaging; SD: standard deviation.

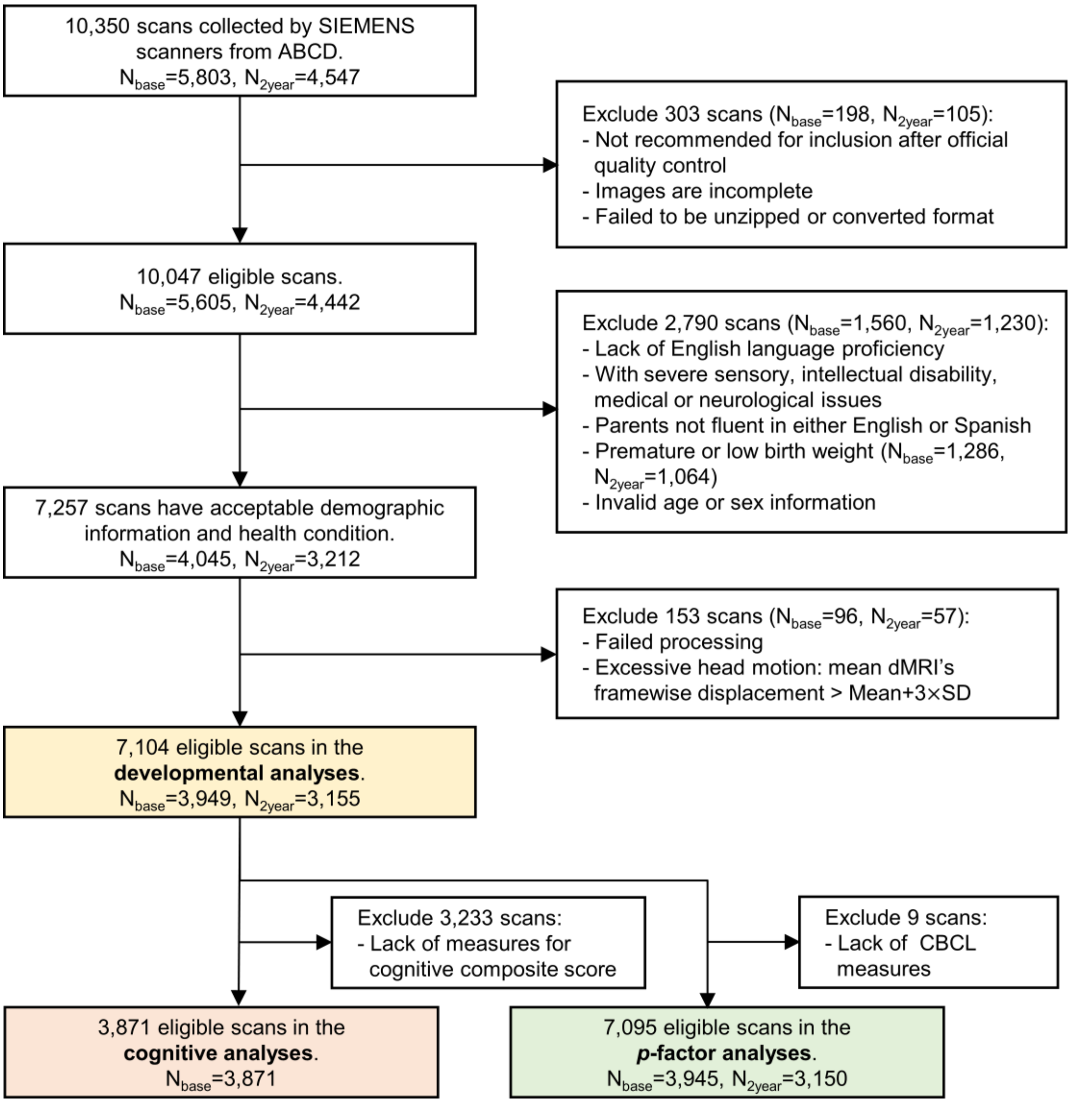


**Fig. S2. Flowchart of inclusion and exclusion for participants in the ABCD dataset.** ABCD: the Adolescent Brain Cognitive Development; CBCL: Child Behavior Checklist.

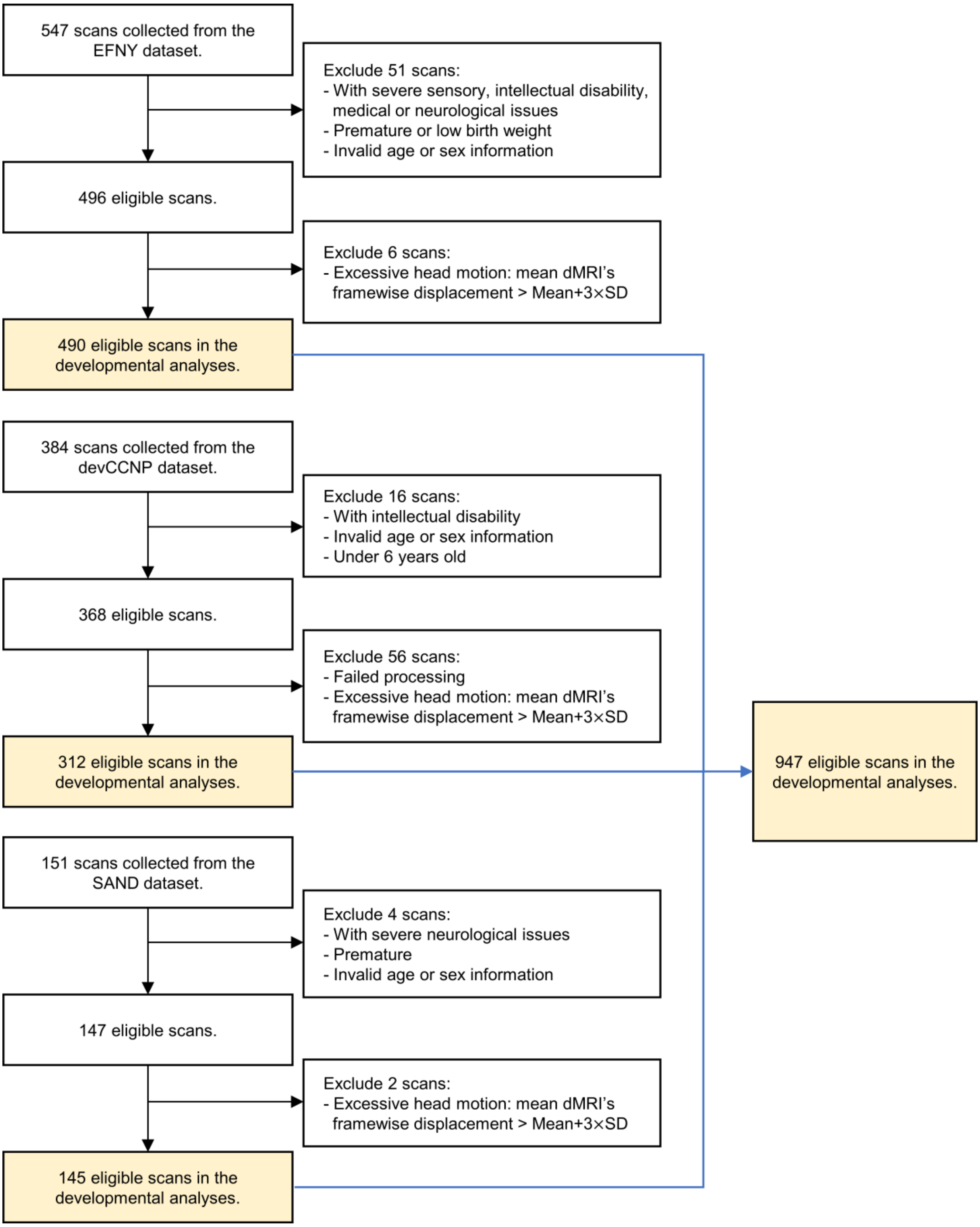


**Fig. S3. Flowchart of inclusion and exclusion for participants in the Chinese Cohort dataset.** devCCNP: the developmental component of the Chinese Color Nest Project; EFNY: the Executive Function and Neurodevelopment in Youth; SAND: Shandong Adolescent Neuroimaging Project on Depression.

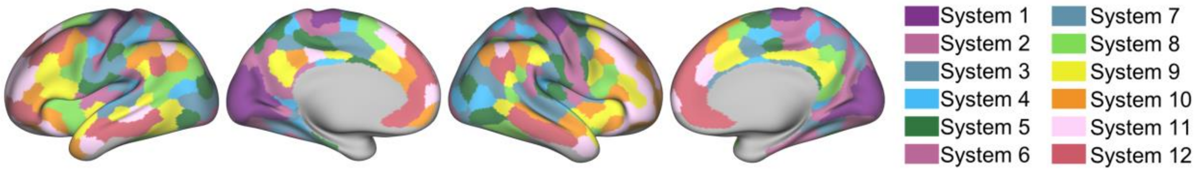


**Fig. S4. Discrete visualization of the 12 cortical systems defined by partitioning the S-A axis.**

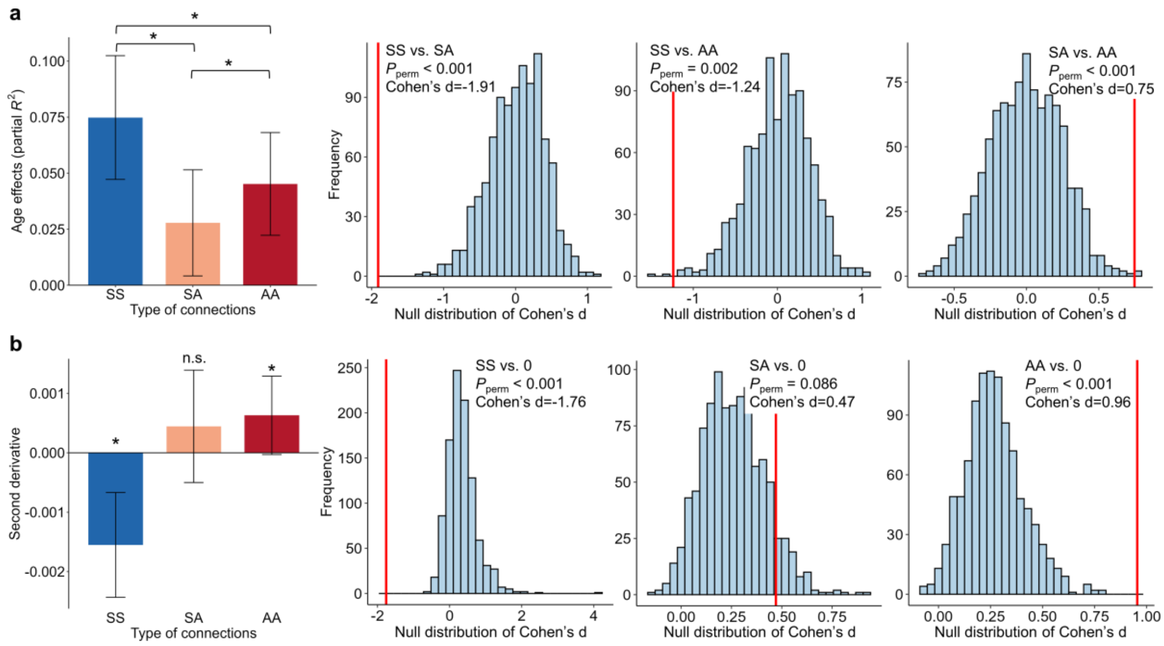


**Fig. S5. Comparison of developmental features across different types of connections. a**, The 12 systems were classified into two categories based on their spatial overlap with the Yeo-7 networks. The four systems closest to the sensorimotor pole were assigned to the sensorimotor group ("S"), and the remaining eight systems were assigned to the association group ("A"). Accordingly, 78 connections were categorized into three types, SS, SA, and AA. The mean age effects (partial $R^2$) and corresponding standard deviations are shown for each connection type. Pairwise differences were quantified using Cohen's d. Null distributions were generated by permutation testing (1,000 permutations; light blue), and observed Cohen's d values are shown as red vertical lines. All pairwise differences were significant. **b**, Mean second derivatives and corresponding standard deviations are shown for each connection type, and differences between each connection type and zero were quantified using Cohen's d. Permutation tests (1,000 permutations) indicated that second-derivative values for SS connections were significantly lower than zero, whereas those for AA connections were significantly greater than zero. SS, sensorimotor-sensorimotor; SA, sensorimotor-association; AA, association-association; *: $P_{perm}$ < 0.05; n.s.: not significant.

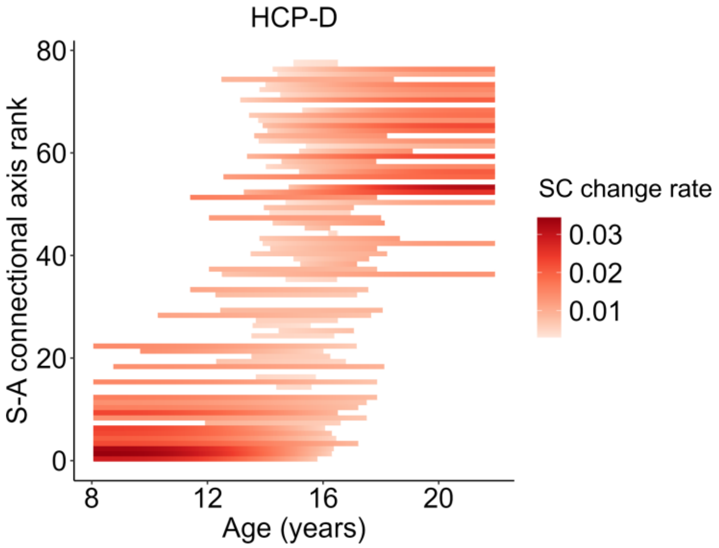


**Fig. S6. Significant developmental rates of the large-scale structural connections in the HCP-D dataset**. The rates of developmental changes of the large-scale structural connections were measured using the first derivatives at 1,000 age points evenly sampled from the age range of 8.1 to 21.9 years. Each row represents an edge, with colors coded based on the magnitudes and direction of the significant developmental rates ($P_{FDR} < 0.05$). Insignificant developmental rates are shown in white. Since all significant derivatives are above zero, we used different shades of red to indicate their magnitudes. SC: structural connectivity; S-A: sensorimotor-association.

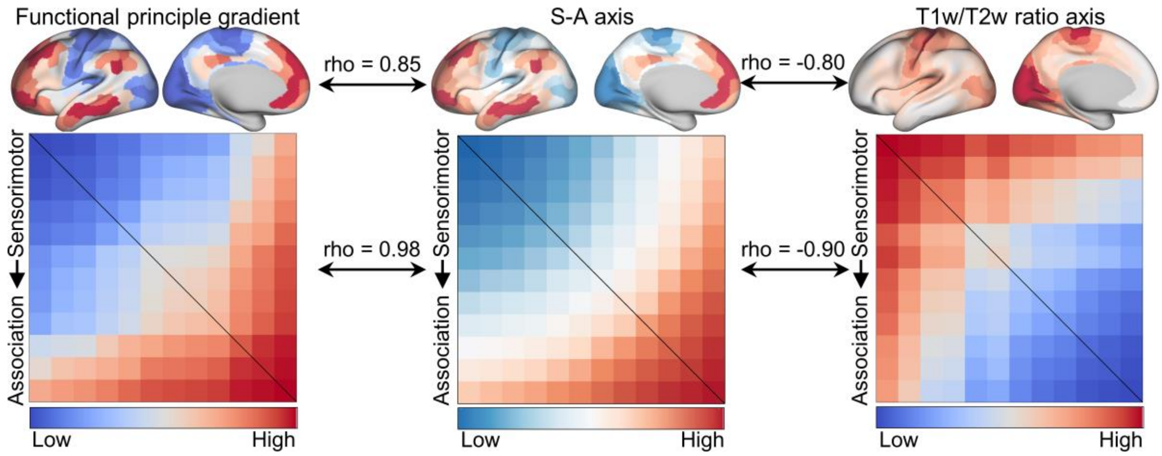


**Fig. S7. Comparison of S-A axis with alternative organizational principles.** Cortical maps of the S-A axis showed a strong positive correlation with the principal functional gradient (rho = 0.85) and a strong negative correlation with the T1w/T2w ratio (rho = -0.80) across cortical vertices. Mean values of the principal functional gradient and T1w/T2w ratio were computed and visualized within each of the 12 cortical systems. These system-level measures were subsequently converted to connectome-level axes using a sum-of-squares approach and rank-ordered to yield discrete integer values from 1 to 78. The S-A connectional axis was highly correlated with both the functional gradient-based connectional axis (rho = 0.98) and the T1w/T2w-based connectional axis (rho = -0.90). S-A: sensorimotor-association.

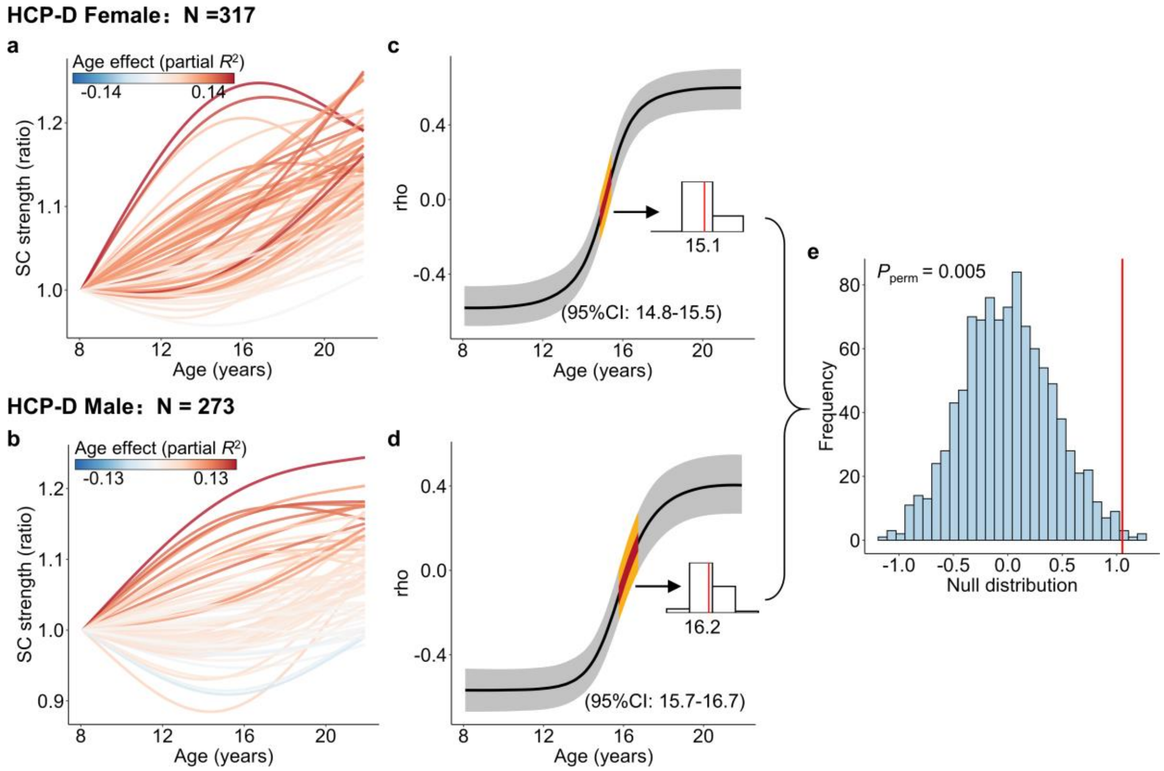


**Fig. S8. Sex-stratified developmental trajectories and alignment with the S-A connectional axis.** **a**, Developmental trajectories of SC strength in females (N = 317). Each line represents one system-to-system connection, color-coded by the magnitude of age effects (partial $R^2$). **b**, Developmental trajectories of SC strength in males (N = 273). **c**, Age-resolved alignment between the spatial pattern of SC developmental rates (first derivatives) and the S-A connectional axis in females, with zero alignment occurring at 15.1 years (95% CI: 14.8–15.5). **d**, Age-resolved alignment between the spatial pattern of SC developmental rates and the S-A connectional axis in males, with zero alignment occurring at 16.2 years (95% CI: 15.7–16.7). **e**, Null distribution of the difference in median zero-crossing age between females and males from 1,000 permutations of sex labels. The red vertical line represents the observed sex difference, with the corresponding permutation-based significance ($P_{perm}$ = 0.005). SC: structural connectivity; CI: credible interval.

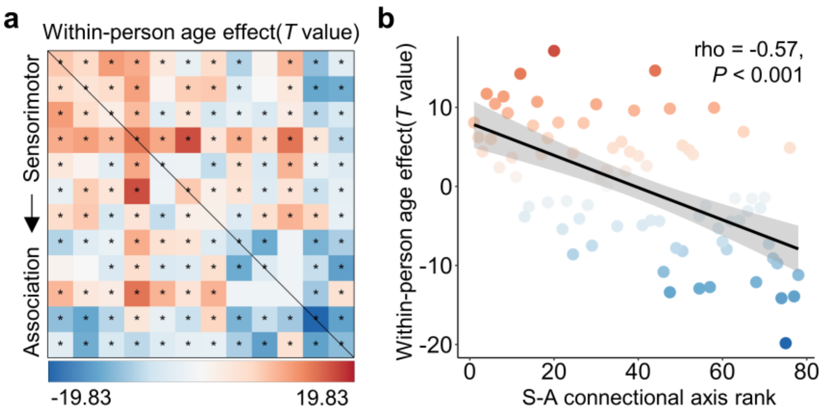


**Fig. S9. Within-person age effects in SC strength were associated with the S-A connectional axis ranks. a**, Connection-wise within-person age effects estimated from within–between linear mixed-effects models in the ABCD dataset. Values represent *T* statistics for the within-person age term across the 78 system-to-system connections with asterisks indicating significant within-person age effects ($P_{FDR} < 0.05$). **b**, Scatter plot showing the relationship between within-person age effects (*T* values) and S-A connectional axis ranks across the 78 connections. S-A: sensorimotor-association.

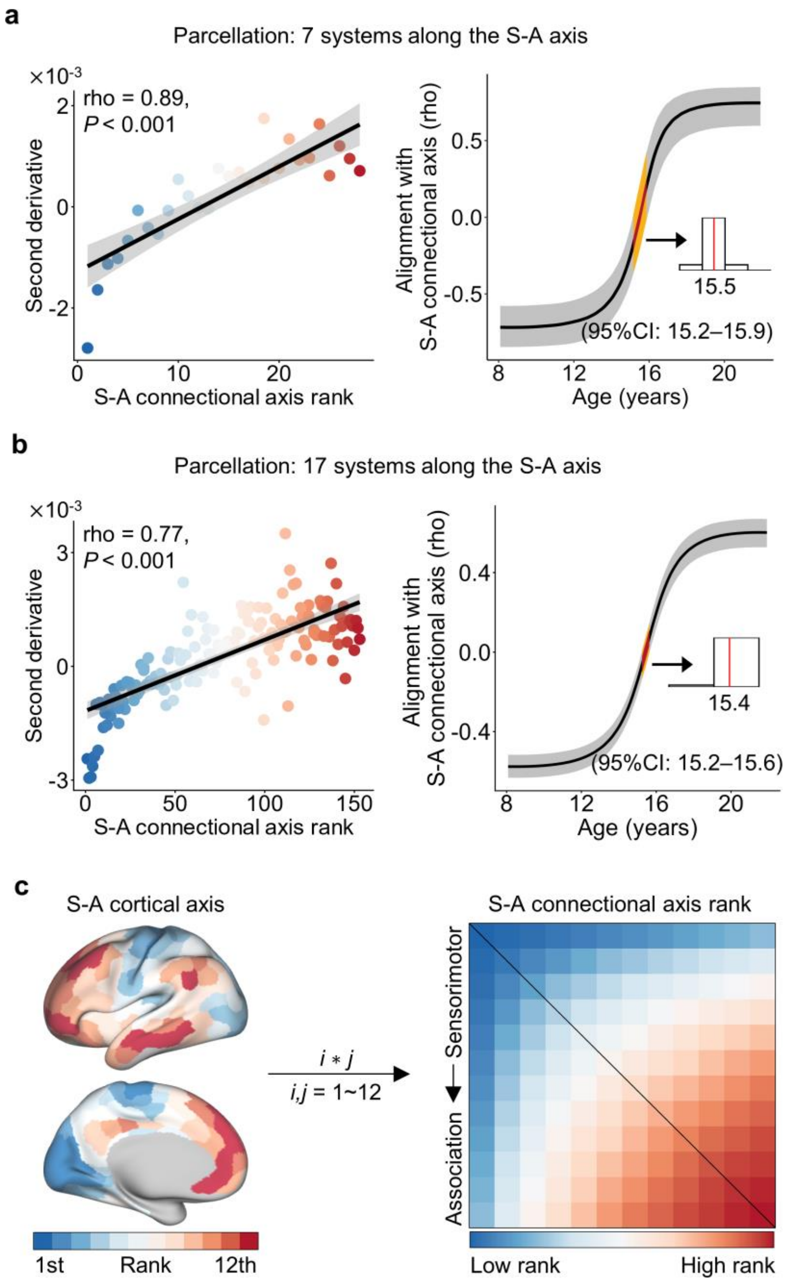


**Fig. S10. Sensitivity analyses using alternative network scales and an alternative S-A connectional axis definition.** We first replicated the key results using structural connectomes with alternative network scales of 7 (**a**) and 17 (**b**). For each scale, the left panel shows the second derivatives of developmental trajectories are strongly correlated with the S-A connectional axis ranks. The right panel shows the age-resolved alignment between the spatial pattern of structural-connectivity development slopes and the S-A axis; the yellow band marks the age window of zero alignment, with the median age annotated. **c**, We additionally defined an alternative S-A connectional axis by multiplying the S-A cortical axis ranks of each pair of nodes. Results obtained using this alternative axis are shown in **Fig. 6h**. S-A: sensorimotor-association.

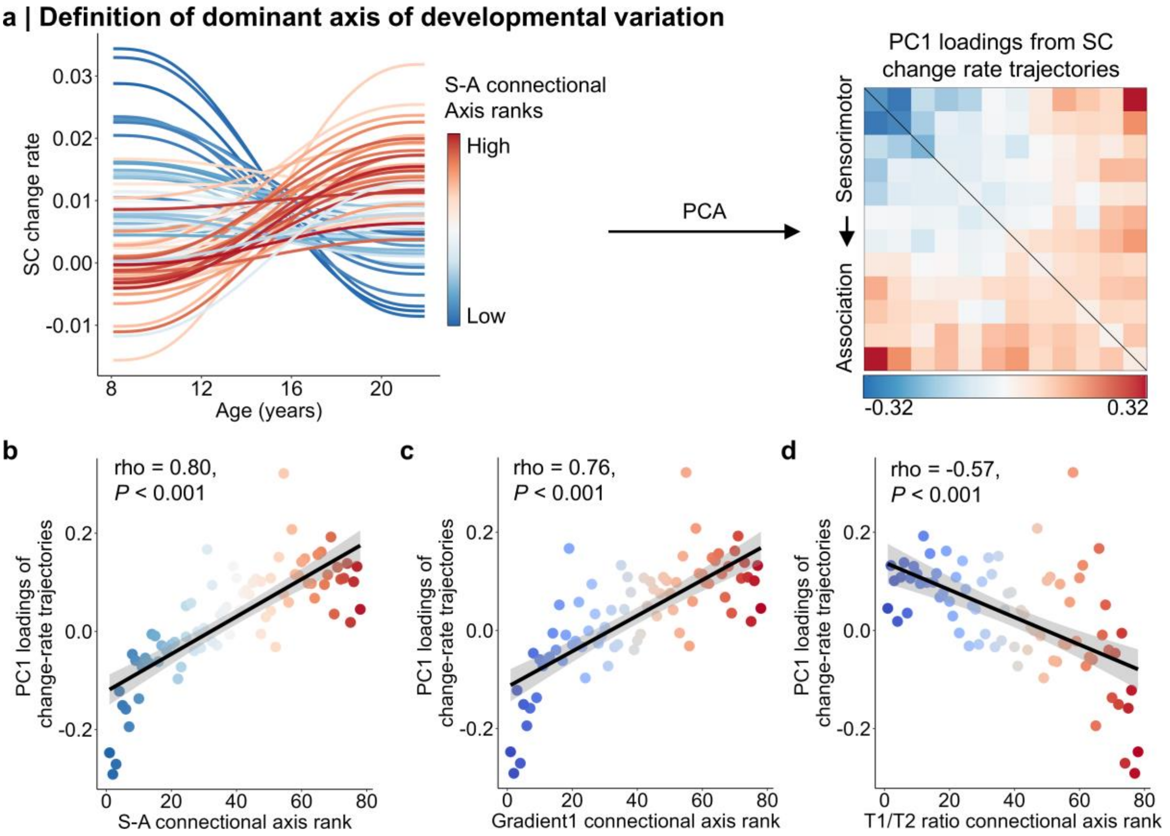


**Fig. S11. Data-driven developmental axis and its alignment with connectional axes. a,** The data-driven dominant developmental axis was defined as the loadings of the first principal component (PC1) obtained from PCA applied to SC change-rate (first-derivative) trajectories. **b**, The data-driven developmental axis was strongly correlated with the S-A connectional axis across all connections (rho = 0.80, $P < 0.001$). **c**, The data-driven developmental axis was strongly associated with the connectional axis derived from the principal functional gradient (rho = 0.76, $P < 0.001$). **d**, The data-driven developmental axis was negatively correlated with the T1w/T2w-based connectional axis (rho = -0.57, $P < 0.001$). SC: structural connectivity; PCA: principal component analysis; PC1: the first principal component; S-A: sensorimotor-association.

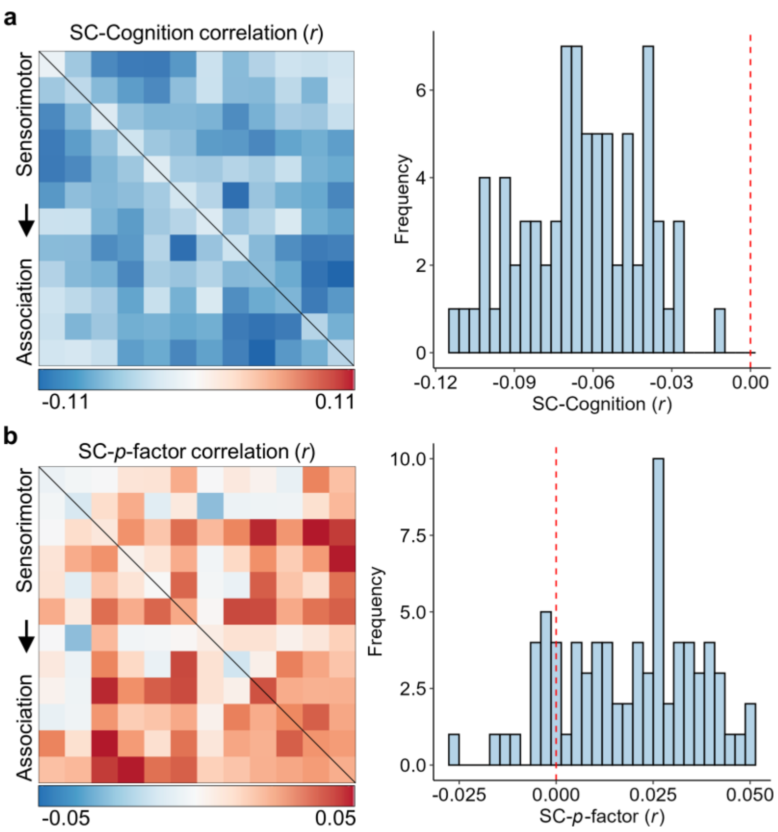


**Fig. S12. Partial correlation coefficients of SC–cognition and SC–*p*-factor associations. a**, Partial correlation coefficients between SC strength and fluid composite cognition scores, controlling for age, sex, and head motion. Correlation coefficients ranged from -0.11 to -0.01. **b**, Partial correlation coefficients between SC strength and *p*-factor scores, controlling for age, sex, and head motion. Correlation coefficients ranged from -0.03 to 0.05. SC: structural connectivity.

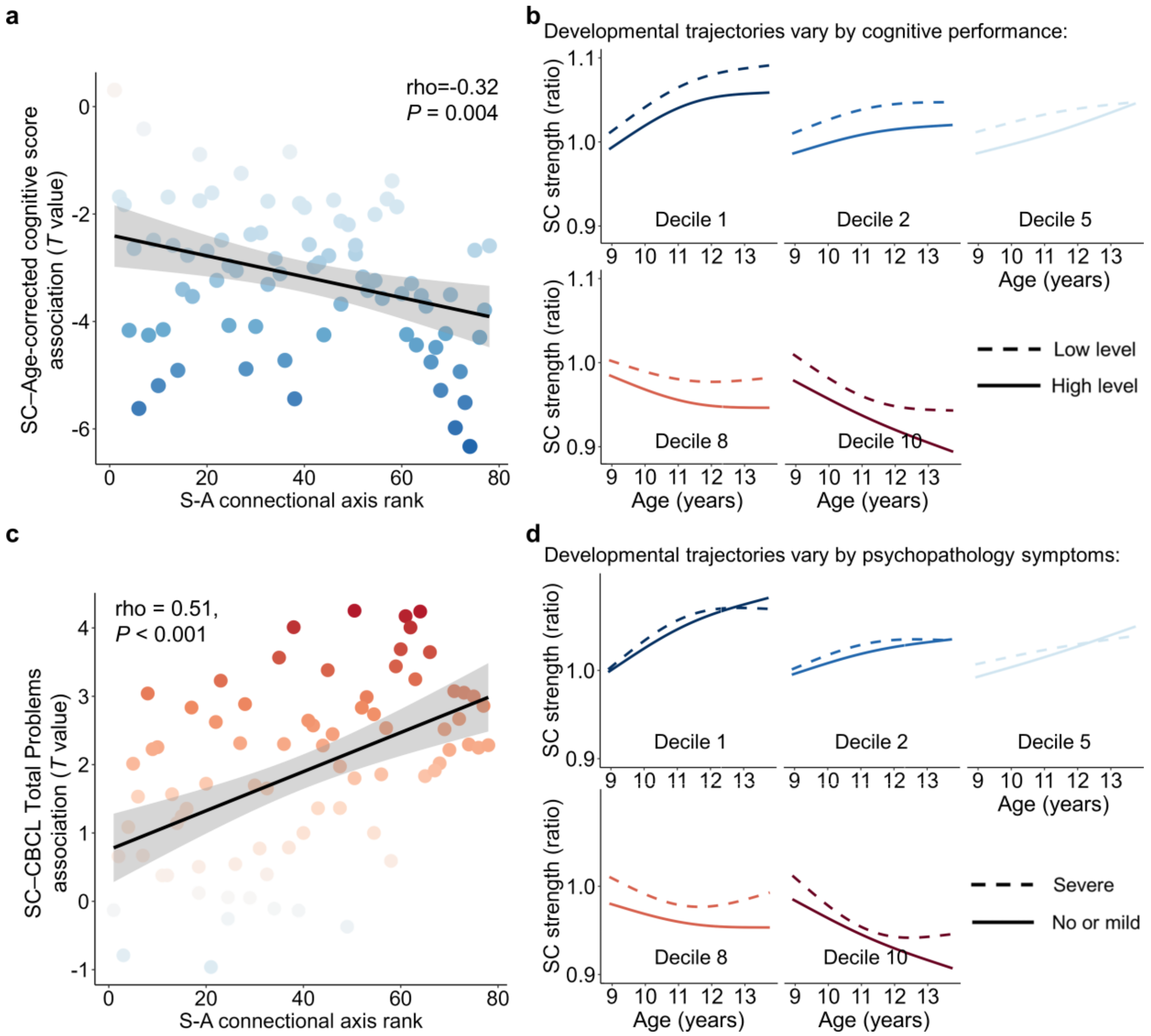


**Fig. S13. Sensitivity analyses using alternative measures of cognition and psychopathology.** **a**, Across all connections, the effect sizes of SC–cognition associations were significantly correlated with S-A connectional axis ranks (rho = -0.32, $P = 0.004$) when using age-corrected NIH Toolbox fluid composite scores. **b**, Developmental trajectories of SC strength are shown for groups with low (10th percentile) and high (90th percentile) baseline cognitive performance across five deciles of the S-A connectional axis. **c**, The spatial distribution of SC–psychopathology associations was significantly aligned with the S-A connectional axis (rho = 0.51, $P < 0.001$) when psychopathology was indexed using the CBCL Total Problems score. **d**, Developmental trajectories of SC strength are shown at low (10th percentile) and high (90th percentile) CBCL Total Problems scores across five deciles of the S-A connectional axis. CBCL: Child Behavior Checklist; SC: structural connectivity.

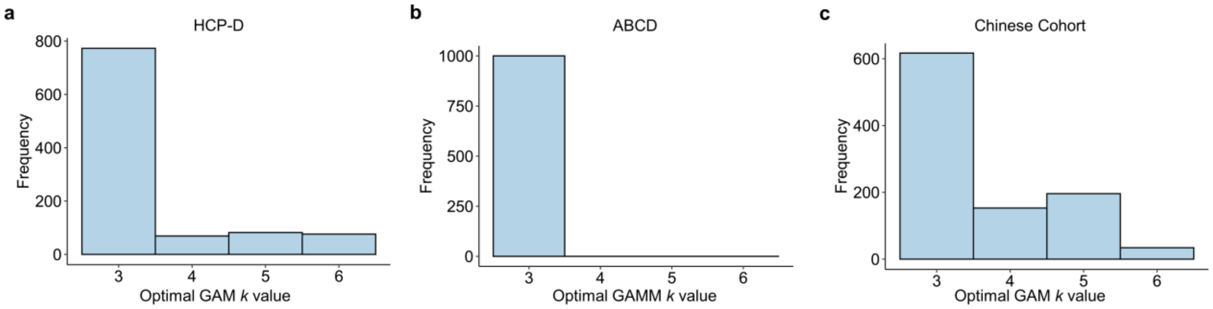


**Fig. S14. Selection frequency of *k* values across bootstrap iterations.** Across 1,000 iterations, a *k* value of 3 was selected as the optimal choice in 773 iterations for the HCP-D dataset (**a**), 1,000 iterations for the ABCD dataset (**b**), 617 iterations for the Chinese Cohort dataset (**c**).

**Table S1. Demographic and cognitive characteristics of participants from the HCP-D dataset.**

| | HCP-D dataset |
|---|---|
| N | 590 |
| Age (years) (mean (SD)) | 14.72 (3.92) |
| Sex = Male (%) | 273 (46.3) |
| Handedness (%) | |
| Right-handed | 519 (88.0) |
| Left-handed | 43 (7.3) |
| Mixed handed | 28 (4.7) |
| Race/ethnicity (%) | |
| Hispanic | 86 (14.6) |
| Non-Hispanic Asian | 43 (7.5) |
| Non-Hispanic Black | 56 (9.5) |
| Non-Hispanic White | 344 (58.3) |
| Others | 61 (10.3) |
| Average cognitive performance (mean (SD)) | 107.76 (12.75) |
| Mean FD (mean (SD)) | 0.66 (0.20) |
| Sites (%) | |
| Harvard | 198 (33.6) |
| UMinn | 156 (26.4) |
| UCLA | 105 (17.8) |
| WashU | 131 (22.2) |
| Intracranial volume ($mm^3$) (mean (SD)) | 1598201.51 (154010.77) |
| Family income-to-needs ratio | 5.48 (6.04) |

Note: The number of participants and percentages were displayed for categorical variables, while mean and standard deviation (SD) were provided for numeric variables. Ages in months and sex were extracted from 'HCD_LS_2.0_subject_completeness.csv' in the HCP-D dataset. Ages were converted to years by dividing by 12. Handedness scores derived from 'edinburgh_hand01.txt' were converted into a 3-level factor. Scores over 60 were defined as right-handed, under -60 as left-handed, and those in the middle as mixed-handed[6]. Race was obtained from 'socdem01.txt'. Average cognitive performance without age correction were derived from 'cogcomp01.txt'. Mean framewise displacement (FD) measures head motion during diffusion MRI scanning[7]. Site information was derived from 'ndar_subject01.txt'. The family income-to-needs ratio was calculated by dividing the annual family income ('socdem01.txt') by the federal poverty line for the year of the interview and the family size.

**Table S2. Demographic, cognitive, and psychiatric characteristics of participants in the ABCD dataset.**

| | Baseline | Two-year follow-up |
|---|---|---|
| N | 3949 | 3155 |
| Age (years) (mean (SD)) | 9.93 (0.63) | 11.95 (0.65) |
| Sex = male (%) | 2075 (52.5) | 1701 (53.9) |
| Race/ethnicity (%) | | |
| Hispanic | 678 (17.2) | 525 (16.6) |
| Non-Hispanic Asian | 68 (1.7) | 41 (1.3) |
| Non-Hispanic Black | 575 (14.6) | 476 (15.1) |
| Non-Hispanic White | 2257 (57.2) | 1817 (57.6) |
| Other | 371 (9.4) | 296 (9.4) |
| Handedness (%) | | |
| Right-handed | 3166(80.2) | 2510(79.6) |
| Left-handed | 270(6.8) | 230(7.3) |
| Mixed handed | 513(13.0) | 415(13.2) |
| Average cognitive performance (mean (SD)) | 92.53 (10.12) | - |
| *P*-factor (mean (SD)) | 0.07(0.83) | 0.04(0.82) |
| Mean FD (mean (SD)) | 0.55 (0.21) | 0.53 (0.20) |
| Sites (%) | | |
| Site02 | 174(4.4) | 161(5.1) |
| Site03 | 308(7.8) | 248(7.9) |
| Site05 | 225(5.7) | 183(5.8) |
| Site06 | 392(9.9) | 314(10.0) |
| Site07 | 153(3.9) | 123(3.9) |
| Site09 | 258(6.5) | 162(5.1) |
| Site11 | 268(6.8) | 190(6.0) |
| Site12 | 444(11.2) | 317(10.0) |
| Site14 | 241(6.1) | 176(5.6) |
| Site15 | 204(5.2) | 168(5.3) |
| Site16 | 696(17.6) | 581(18.4) |
| Site20 | 219(5.5) | 226(7.2) |
| Site21 | 367(9.3) | 306(9.7) |
| Intracranial volume ($mm^3$) (mean (SD)) | 1534157.62(132994.37) | 1561853.71(140683.69) |
| Family income-to-needs ratio | 3.80(2.35) | 3.81(2.29) |

Note: The number of participants and percentages were displayed for categorical variables, while mean and SD were displayed for numeric variables. Ages in months and site information were obtained from 'abcd_y_lt.csv'. Ages were then converted to years by dividing by 12. Sex and race/ethnicity were extracted from 'abcd_p_demo.csv'. Three-factor handedness was acquired from 'nc_y_ehis.csv'. Average cognitive performance without age correction were obtained from 'nc_y_nihtb.csv'. General psychopathology factor (*p*-factor) scores were derived from the CBCL. The family income-to-needs ratio was calculated by dividing the annual family income (‘abcd_p_demo.csv’) by the federal poverty line for the year of the interview and the family size.

**Table S3. Demographic and cognitive characteristics of participants from the Chinese Cohort dataset.**

| | devCCNP-ses1 | devCCNP-ses2 | devCCNP-ses3 | EFNY | SAND |
|---|---|---|---|---|---|
| N | 210 | 73 | 29 | 490 | 145 |
| Age (years) (mean (SD)) | 9.68 (2.60) | 10.85 (2.34) | 12.17 (2.10) | 14.75 (5.25) | 15.54 (3.24) |
| Sex = male (%) | 120 (57.1) | 41 (56.2) | 20 (69.0) | 249 (50.8) | 51 (35.2) |
| Race (%) | | | | | |
| Asian | 210 (100.0) | 73 (100.0) | 29 (100.0) | 490 (100.0) | 145 (100.0) |
| Handedness (%) | | | | | |
| Right-handed | 184 (93.4) | 70 (97.2) | 29 (100.0) | 279 (68.6) | 101 (78.9) |
| Left-handed | 5 (2.5) | 1 (1.4) | 0 (0.0) | 7 (1.7) | 0 (0.0) |
| Mixed handed | 8 (4.1) | 1 (1.4) | 0 (0.0) | 121 (29.7) | 27 (21.1) |
| Mean FD (mean (SD)) | 0.59 (0.34) | 0.49 (0.27) | 0.41 (0.17) | 0.56 (0.11) | 0.59 (0.19) |
| Intracranial volume ($mm^3$) (mean (SD)) | 1450508.93 (134656.16) | 1493327.38 (132939.62) | 1590492.17 (157471.04) | 1527375.03 (159414.76) | 1427493.68 (216870.43) |

Note: The number of participants and percentages were displayed for categorical variables, while mean and SD were provided for numeric variables. The demographic information from the devCCNP was obtained from the participant information table. The demographic information of the EFNY and SAND was provided by Z.C., Y.L. and K.W.

**Table S4. Image acquisition parameters for T1-weighted images and diffusion MRI for each dataset.**

| | Sequence | TR (ms) | TE (ms) | TI (ms) | Flip angle (°) | FOV ($mm^2$) | Slices | Voxel Size (mm) | Diffusion directions | b-values ($s/mm^2$) | Acquisition time (min:s) |
|---|---|---|---|---|---|---|---|---|---|---|---|
| HCP-D (3T SIEMENS Prisma with 32-channel head coil) | | | | | | | | | | | |
| T1WI | MPRAGE | 2500 | 1.8,3.6, 5.4,7.2 | 1000 | 8 | 256×256 | 208 | 0.8 | NA | NA | 8:22 |
| dMRI (AP/PA) | Multiband EPI | 3230 | 89.2 | NA | 78 | 210×210 | 92 | 1.5 | 185 | 1500, 3000 | 5:37*2runs* 2sessions |
| ABCD (3T SIEMENS) | | | | | | | | | | | |
| T1WI | MPRAGE | 2500 | 2.88 | 1060 | 8 | 256×256 | 176 | 1.0 | NA | NA | 7:12 |
| dMRI (PA) | Multiband EPI | 4100 | 88 | NA | 90 | 240×240 | 81 | 1.7 | 96 | 500, 1000, 2000, 3000 | 7:31 |
| fmap (AP) | Single-band EPI | 12400 | 89 | NA | 90 | 240×240 | 81 | 1.7 | 0 | 0 | Not available |
| Chinese Cohort-EFNY (3T SIEMENS Prisma with 64-channel head coil) | | | | | | | | | | | |
| T1WI | MPRAGE | 1500 | 1.87 | 756 | 10 | 256×256 | 208 | 0.8 | NA | NA | 7:15 |
| dMRI (PA) | Multiband EPI | 3100 | 86 | NA | 90 | 205×205 | 84 | 1.8 | 120 | 500, 1000, 2000, 3000 | 7:00 |
| fmap (AP) | Multiband EPI | 3100 | 86 | NA | 90 | 205×205 | 84 | 1.8 | 0 | 0 | 0:32 |
| Chinese Cohort-devCCNP (3T GE discovery MR750 with 8-channel head coil) | | | | | | | | | | | |
| T1WI | 3D SPGR | 6.7 | 2.9 | 450 | 12 | 256×256 | 176 | 1.0 | NA | NA | 4:41 |
| dMRI (PA) | Single-band EPI | 8724 | 81.4 | NA | 90 | 224×224 | 75 | 2.0 | 64 | 1000 | 10:54 |

**Table S4 (continued). Image acquisition parameters for T1-weighted images and diffusion MRI for each dataset.**

| | Sequence | TR (ms) | TE (ms) | TI (ms) | Flip angle (°) | FOV ($mm^2$) | Slices | Voxel Size (mm) | Diffusion directions | b-values ($s/mm^2$) | Acquisition time (min:s) |
|---|---|---|---|---|---|---|---|---|---|---|---|
| Chinese Cohort-SAND (3T SIEMENS Verio) | | | | | | | | | | | |
| T1WI | MPRAGE | 2400 | 2.2 | 1000 | 8 | 256×256 | 208 | 0.9 | NA | NA | Not available |
| dMRI (AP) | Single-band EPI | 9635 | 100 | NA | 90 | 256×256 | 66 | 2 | 64 | 1000 | Not available |

Note: T1WI: T1-weighted imaging; dMRI: diffusion magnetic resonance imaging; MPRAGE: magnetization prepared rapid gradient echo; EPI: echo planar imaging; 3D SPGR: three-dimensional spoiled gradient recalled; AP: anterior-to-posterior phase-encoding direction; PA: posterior-to-anterior phase-encoding direction.
